# A Substrate-triggered µ-Peroxodiiron(III) Intermediate in the 4-Chloro-L-Lysine-Fragmenting Heme-Oxygenase-like Diiron Oxidase (HDO) BesC: Substrate Dissociation from, and C4 Targeting by, the Intermediate

**DOI:** 10.1101/2021.12.02.471016

**Authors:** Molly J. McBride, Mrutyunjay A. Nair, Debangsu Sil, Jeffrey W. Slater, Monica Neugebauer, Michelle C. Y. Chang, Amie K. Boal, Carsten Krebs, J. Martin Bollinger

**Affiliations:** Department of Chemistry, The Pennsylvania State University, University Park, Pennsylvania 16802, United States; Department of Biochemistry and Molecular Biology, The Pennsylvania State University, University Park, Pennsylvania 16802, United States; Department of Chemical and Biomolecular Engineering, University of California, Berkeley, Berkeley, CA, USA; Departments of Chemistry and of Molecular and Cell Biology, University of California, Berkeley, and Lawrence Berkeley National Laboratory, Berkeley, CA, USA; Department of Systems Biology, Harvard Medical School, Boston, Massachusetts 02115, United States

## Abstract

The enzyme BesC from the *<u>β</u>*-<u>e</u>thynyl-L-<u>s</u>erine biosynthetic pathway in *Streptomyces cattleya* fragments 4-chloro-L-lysine (produced from L-Lysine by BesD) to ammonia, formaldehyde, and 4-chloro-L-allylglycine and can analogously fragment L-Lys itself. BesC belongs to the emerging family of O_2_-activating non-heme-diiron enzymes with the “heme-oxygenase-like” protein fold (HDOs). Here we show that binding of L-Lys or an analog triggers capture of O_2_ by the protein’s diiron(II) cofactor to form a blue µ-peroxodiiron(III) intermediate analogous to those previously characterized in two other HDOs, the olefin-installing fatty acid decarboxylase, UndA, and the guanidino-*N*-oxygenase domain of SznF. The ∼ 5- and ∼ 30-fold faster decay of the intermediate in reactions with 4-thia-L-Lys and (4*RS*)-chloro-DL-lysine than in the reaction with L-Lys itself, and the primary deuterium kinetic isotope effects (D-KIEs) on decay of the intermediate and production of L-allylglycine in the reaction with 4,4,5,5-[^2^H]-L-Lys, imply that the peroxide intermediate or a successor complex with which it reversibly interconverts initiates the oxidative fragmentation by abstracting hydrogen from C4. Surprisingly, the sluggish substrate L-Lys can dissociate after triggering the intermediate to form, thereby allowing one of the better substrates to bind and react. Observed linkage between Fe(II) and substrate binding suggests that the triggering event involves a previously documented (in SznF) ordering of the dynamic HDO architecture that contributes one of the iron sites, a hypothesis consistent with the observation that the diiron(III) product cluster produced upon decay of the intermediate spontaneously degrades, as it has been shown to do in all other HDOs studied to date.

## INTRODUCTION

Decades of study of ferritin-like non-heme diiron oxygenases and oxidases (FDOs) established a functional paradigm for these enzymes^1–2^ involving oxidative addition of O_2_ to the fully reduced form of the His_2-3_(Glu/Asp)_4_-coordinated cofactor to form a µ-(hydro)peroxodiiron(III) complex that either directly oxidizes the substrate by oxygen-atom transfer (OAT)^3–5^ or undergoes O-O-bond cleavage (reductively or without redox) to a high-valent [Fe_2_(III/IV) or Fe_2_(IV/IV)] intermediate that then initiates the relevant oxidation by abstracting hydrogen (hydrogen atom transfer, HAT).^6–11^ In many FDOs, binding of the substrate or an effector protein is required to “trigger” formation of the O_2_-reactive configuration of the cofactor by, for example, inducing a carboxylate shift.^12–13^ With the exception of one of two reactions mediated by each of a pair of arylamine *N*-oxygenases, AurF and CmlI,^5, 14^ completion of the relevant oxidation leaves a stable diiron(III) “product” form of the cluster that must undergo *in situ* reduction by a general electron-transfer protein (e.g., ferredoxin) or a specific reductase to reset the cofactor for the next turnover.^1^

Most FDOs bind iron stably, and multiple members of the family have been isolated with their diiron cofactors intact.^4–5, 15–17^ By contrast, the first few characterized members of the emerging new family of nonheme-diiron oxidases and oxygenases sharing the heme-oxygenase-like protein fold (HDOs) were all initially characterized with, at most, one of the two iron ions of the cofactor bound.^18–21^ Indeed, the founding member of the HDO family, the olefin-forming fatty acid decarboxylase, UndA, was initially thought to use a mono-iron cofactor, because it was isolated and structurally characterized with iron bound only in what is now conventionally denoted subsite 1 of the diiron site.^18^ More recent biochemical and biophysical analysis of UndA showed that the functional iron cofactor is, as in the FDOs, dinuclear.^22–23^ One of these studies reported rapid accumulation and decay of a µ-peroxodiiron(III) intermediate upon mixing of the UndA•Fe(II)_2_•dodecanoic acid complex with O_2_-saturated buffer and further demonstrated the slow (minutes-hours) disintegration of the product diiron(III) cluster, thus rationalizing the prior observation of the mono-iron form.^23^ Subsequent analysis of the tri-domain enzyme, SznF, which produces the nitrosourea pharmacophore of the anticancer drug, streptozotocin, mirrored these observations on UndA.^24–25^ The central HDO domain of SznF catalyzes sequential OATs to *N^δ^* of *N^ω^*-methyl-L-arginine and then to N*^ω^*^’^ of the hydroxylated intermediate before the enzyme’s C-terminal mono-iron cupin domain oxidatively rearranges the dihydroxy compound to *N^ω^*-methyl-*N^w^*-nitroso-L-citrulline.^19^ A µ-peroxodiiron(III) complex was shown to accumulate upon exposure of the SznF•Fe(II)_2_ complex to O_2_ and to react efficiently with either *N^ω^*-methyl-L-arginine or the *N^δ^*-hydroxylated intermediate, and the product diiron(III) cluster was again shown to disintegrate on the minutes-to-hours timescale.^24^ Interestingly, substrate triggering of intermediate formation was not seen for SznF, by contrast to the case of UndA. This dichotomy within the first two mechanistically characterized HDOs mirrors that seen in the FDOs, of which the *N*-oxygenases AurF and CmlI were shown not to require triggering by either the substrate or an accessory protein.^24^

Recent x-ray crystallographic studies of SznF^26^ shed light on the instability of the diiron cofactor in its HDO domain,^19^ and structural characteristics shared with UndA^18, 23^ suggested that the insight might be general to other (potentially all) enzymes in the emerging HDO family.^1^ A strategy of (i) preparing naive apo protein by heterologous expression in low-iron medium supplemented with Mn(II) to occupy the metal sites and prevent potentially deleterious cofactor cycling during protein production, (ii) chelation of the bound Mn(II) from the purified protein, (iii) crystallization of the apo protein, and (iv) soaking of crystals with Fe(II) yielded a structure of the SznF•Fe(II)_2_ complex with both sites almost fully occupied, the first such structure of an HDO.^26^ Comparison of apo and metallated structures revealed that one of the iron sites in the HDO domain (site 2) is not pre-formed (Fig. S1), by contrast to observations made on structures of apo FDOs.^27–30^ An 8-residue segment of the protein was seen to rearrange from a series of turns in the apo state to a canonical *α*-helix (part of *α*3) in the diiron(II) complex, in the process relocating a site 2 carboxylate ligand (D315) ∼10 Å from the protein surface into the helical core.^26^ It was posited that dynamics of the modestly stable *α*3 could initiate the observed disintegration of the product diiron(III) complex. Given that a segment of *α*3 in UndA could not even be confidently modeled because of poor (or absent) electron density,^23^ this explanation for cofactor instability could be applicable also to the founding HDO. The structure of diiron(II) SznF also revealed an unexpected, additional carboxylate ligand that gives the iron in site 1 four protein-derived ligands (His_2_-carboxylate_2_) in its first coordination sphere.^26^ This extra carboxylate is not present in UndA (nor conserved in the majority of other hypothetical HDOs identified bioinformatically) but is conserved in the 2-aminoimidazole *N*-oxygenase RohS from the azomycin biosynthetic pathway,^31^ leading to the hypothesis that the presence of the ligand might be an identifying trait of heteroatom oxygenases in the HDO family. Moreover, the fact that, in UndA, the *substrate* carboxylate ligates Fe1 at essentially the same site^23^ led to the suggestion that such an interaction might contribute to substrate triggering in HDOs lacking the extra site 1 carboxylate.

A recent mapping of the ***Streptomyces cattleya*** biosynthetic pathway to the acetylenic amino acid *<u>β</u>*-<u>e</u>thynyl-L-<u>s</u>erine showed that the iron(II)- and 2-oxoglutarate-dependent aliphatic halogenase, BesD, installs chlorine upon C4 of L-lysine and the bioinformatically assigned HDO, BesC, then fragments the 4-chloro-L-lysine (4-Cl-L-Lys) to ammonia, formaldehyde, and 4-chloro-L-allylglycine, which is further processed to the namesake product by chloride elimination and C3 hydroxylation (Scheme 1).^21, 32^ This same study reported that the 4-Cl substituent installed by BesD is not strictly required by BesC: L-Lys itself is also fragmented to unsubstituted L-allylglycine.^21^ Both reactions mediated by BesC cleave C4-H, C5-C6, and C6-N7 bonds of the substrate in a net two-electron oxidation requiring dioxygen. Multiple mechanisms can be envisaged for this reaction (Scheme S1); a subset have hydrogen-atom transfer (HAT) from C4 to a diiron(II)- and O_2_-derived intermediate as the initiating step (Scheme S1B). As for UndA and SznF,^18–19^ stable binding of a diiron cluster was not demonstrated in the initial study of BesC.^21^

**Scheme 1.**
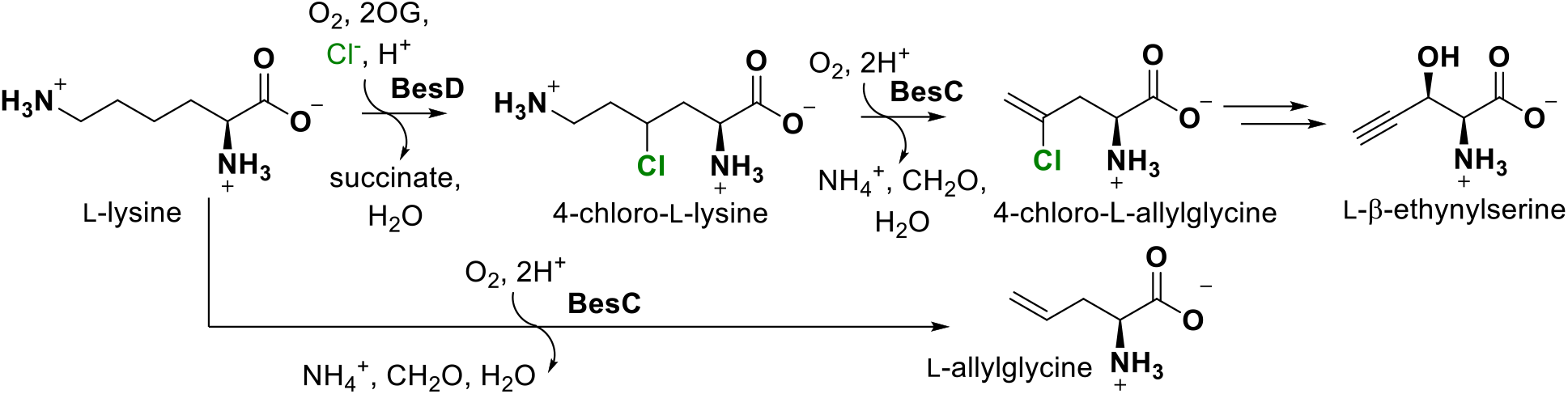
The bes biosynthetic pathway involving iron(II)- and 2-oxoglutarate-dependent halogenase, BesD, and desaturase/lyase, BesC. The lower path indicates that BesC also analogously fragments L-lysine.

Here we show that, in common with UndA but in contrast to SznF, BesC is profoundly substrate triggered in its reaction with dioxygen. In the presence of L-lysine or an analog, but not in its absence, the enzyme’s diiron(II) complex can efficiently capture O_2_ in a relatively long-lived µ-peroxodiiron(III) complex spectroscopically similar to those recently characterized in the other two HDOs.^23–24^ Triggering occurs by the substrate-driven proper configuration of an initially non-functional iron site, which, given prior observations on the other HDOs, is most likely site 2. Remarkably, once the substrate has promoted diiron(II) cofactor assembly and O_2_ capture, it can then dissociate. The observations of substrate-driven cofactor assembly, substrate dissociation from the reactive complex, and disintegration of the diiron(III) product cluster in BesC again underscore the dynamic nature of the HDO architecture in its acquisition and release of reaction components, as noted in previous studies on UndA and SznF.^23–24^ Moreover, the markedly faster decay of the complex in the presence of L-Lys analogs modified at C4 (by H *→* Cl or C *→* S) and the normal primary C4 deuterium kinetic isotope effect (D-KIE) on its decay in the reaction with L-lysine imply that the complex (i) is an authentic intermediate and (ii) either is, or reversibly interconverts with, the species that abstracts hydrogen from C4 to initiate the oxidative fragmentation.

## MATERIALS AND METHODS

### Materials

Isopropyl-β-D-thiogalactopyranoside (IPTG), anhydrous sodium phosphate, and dibasic potassium phosphate were purchased from DOT Scientific. The buffer 4-(2-hydroxyethyl)-1-piperazineethanesulfonic acid (HEPES), glycerol, and imidazole were purchased from Fisher Scientific. Kanamycin was purchased from Teknova. Magnesium sulfate heptahydrate, L-*β*-homolysine and D-Lysine were purchased from Chem-Impex International. Ni(II)-nitrilotriacetic acid agarose (Ni-NTA) resin was purchased from Qiagen. L-Allylglycine was purchased from Cayman Chemical Company. 4,4,5,5-[^2^H_2_]-L-Lysine (*d*_4_-L-Lys) was purchased from Cambridge Isotope Laboratories. L-norleucine was purchased from Alfa Aesar. *Trans*-4,5-dehydro-DL-Lysine was purchased from Bachem. 6-Hydroxy-L-norleucine was purchased from Acrotein Chembio Inc. 4*RS*-Cl-DL-Lysine was purchased from Akos GmbH (Steinen, Germany). EZNA Plasmid DNA Mini Kit was purchased from Omega Bio-Tek. Oligonucleotide primers were purchased from Integrated DNA Technologies. Polymerase chain reaction (PCR) reagents, Q5 DNA Polymerase Master Mix, restriction enzymes, restriction enzyme buffers, and T4 DNA ligase were purchased from New England Biolabs. *Escherichia coli (Ec)* DH5α and Rosetta(DE3) competent cells were purchased from Novagen. ^57^Fe^0^ was purchased from ISOFLEX, USA. All other chemicals were purchased from Millipore Sigma.

### Preparation of S. cattleya BesC

The coding region from an existing plasmid harboring the gene coding for an N-terminally-His_6_-tagged BesC^21^ was amplified by PCR using the primers shown in Table S1. The PCR product was incorporated into the pET28a(+) vector (Novagen) between its *NdeI* and *Xhol* restriction sites using standard molecular biology protocols. The sequence of the resulting pET28a-BesC construct was verified by Sanger sequencing (Penn State Genomics Core Facility).

The overproduction and purification protocol for BesC was adapted from methods previously described for overexpression of *S. achromogenes* SznF.^24^ The pET28a-BesC vector was used to transform *Ec* Rosetta(DE3) cells. Successful transformants were selected at 37 °C on LB-agar plates supplemented with kanamycin (50 μg/mL). A single colony was used to inoculate a 5 mL Luria broth (LB) starter culture containing 50 µg/mL kanamycin. The starter culture was grown at 37 °C with shaking at 180 rpm for 6 h. The cells were harvested by centrifugation at 10,000*g* for 5 min and resuspended in 5 mL of M9 medium.^33^ A 0.5 mL aliquot of this suspension was used to inoculate a 250 mL M9 starter culture supplemented with 50 μg/mL kanamycin and 50 μM MnCl_2_ along with a trace metals cocktail^34^ (Table S2). The starter culture was incubated at 37 °C overnight (16 h) with shaking at 180 rpm. Aliquots of a starter culture were used to inoculate 1 L cultures of M9 medium supplemented with 50 μg/mL kanamycin and 50 μM MnCl_2_ to an initial OD_600_ of 0.05. The cultures were grown at 37 °C with shaking at 180 rpm to an OD_600_ of ∼ 0.6 ‒ 0.8. After cooling on ice for 30 min with an additional supplement of 200 μM MnCl_2_, BesC overexpression was induced by addition of IPTG to 0.2 mM, and cultures were grown at 18 °C overnight (16-18 h) with shaking at 180 rpm. Cells were harvested by centrifugation at 8000*g* and flash frozen in liquid N_2_. Typically, 8 L of culture yielded 15 g of frozen cell paste.

The frozen cells were resuspended in buffer A [100 mM sodium HEPES,100mM NaCl, 10 mM imidazole, 5% glycerol (*v*/*v*), pH 7.5] supplemented with 80 μg/ml PMSF at a ratio of 5 mL buffer per g of cells. The suspended cells were disrupted via sonication on ice (QSonica 750 W, 20 kHz, 10 s pulse, 30 s off, 60% amplitude, 7 min 30 s total sonication time). The resulting lysate was clarified by centrifugation at 22,000*g*. This step and all subsequent protein purification steps were performed at 4 °C. The supernatant was applied to Ni(II)-nitrilotriacetate (NTA) agarose resin (1 mL resin per 5 mL lysate) and washed with 6 column volumes (CV) of buffer A. Bound proteins were eluted with buffer B [100 mM sodium HEPES, 100mM NaCl 250 mM imidazole, 5% glycerol (*v*/*v*), pH 7.5]. Fractions containing BesC, as determined by SDS-PAGE gel analysis with Coomassie Brilliant Blue staining, were pooled and concentrated in a 10 KDa molecular weight cutoff filtration device (PALL Corporation). To remove adventitiously bound metal ions, the protein was dialyzed overnight into 100 mM sodium HEPES, 100 mM NaCl, 5% (*v*/*v*) glycerol, 10 mM EDTA, pH 7.5. To remove EDTA, the protein was dialyzed against 100 mM sodium HEPES, 100 mM NaCl, 5% (*v*/*v*) glycerol, pH 7.5 for at least 4 h, followed by further separation from small molecules by passage through a pre-packed PD-10 column (GE Healthcare). Protein concentrations were determined by absorbance at 280 nm using a calculated molar extinction coefficient (*ε*_280nm_) of 34,400 M^-1^ cm^-1^ and a predicted molecular weight of 29.4 kDa.^35^ For use in rapid kinetics experiments, protein samples were rendered anoxic on a Schlenk line after purification. Three sets of ten gentle evacuation-refill cycles (with argon gas) were performed, and the anoxic protein was flash frozen and stored in liquid N_2_.

### Rapid Kinetics Experiments

Prior to rapid mixing to initiate reactions, reactant solutions were prepared from anoxic solutions of apo BesC, Fe(II), and substrates in an MBraun anoxic chamber. Stopped-flow absorption experiments were performed at 5 °C in an Applied Photophysics Ltd. (Leatherhead, UK) SX20 stopped-flow spectrophotometer housed in the MBraun chamber and equipped with either a photodiode-array (PDA) or photomultiplier tube (PMT) detector, as previously described.^24^ The path length was 1 cm, except in reactions with ferrozine, in which case it was 0.2 cm. Samples to monitor the BesC reaction by Mössbauer spectroscopy were prepared by the freeze-quench method according to previously published procedures.^36^ Mössbauer spectra were recorded on a spectrometer from SEECO (Edina, MN) equipped with a Janis SVT-400 variable-temperature cryostat. The reported isomer shift is given relative to the centroid of the spectrum of α-iron metal at room temperature. External magnetic fields were applied parallel to direction of propagation of the γ radiation. Simulations of the Mössbauer spectra were carried out using the WMOSS spectral analysis software from SEECO (www.wmoss.org, SEE Co., Edina, MN). More detailed procedures for each experiment can be found in the *Supporting Information* and figure legends.

### Deuterium Kinetic Isotope Effect (D-KIE) on Decay of the µ-Peroxodiiron(III) Complex with L-Lys and 4,4,5,5-[^2^H_4_]-L-Lys

Reactions were carried out with an Agilent 8453 UV-Visible spectrophotometer with a temperature-controlled cuvette holder inside an MBraun anoxic chamber. For reactions carried out at 5 °C, an initial reactant solution of 0.2 mM BesC, 0.4 mM Fe(II), 12 mM L-Lys or 4,4,5,5-[^2^H_4_]-L-Lys (*d*_4_-L-Lys), and 124 mM NaCl in 100 mM sodium HEPES (pH 7.5) and 5% (*v/v*) glycerol was manually mixed with an equal volume of O_2_-saturated buffer (∼1.8 mM). For reactions carried out at 22 °C, a reactant solution of 0.4 mM BesC, 0.8 mM Fe(II), 24 mM L-Lys or *d*_4_-L-Lys, and 148 mM NaCl in 100 mM sodium HEPES (pH 7.5) and 5% glycerol (v/v) was manually mixed with three volume equivalents of O_2_-saturated buffer (∼1.2 mM). Spectra were collected at 5-s intervals for 1000-7200 s after mixing.

### Analysis of BesC Reactions by Liquid Chromatography Mass Spectrometry (LC-MS)

Reactions were initiated by mixing anoxic solutions of BesC, Fe(II) and substrates with O_2_-saturated buffer and terminating the reactions by addition of 1.5 volumes of methanol containing 1% formic acid and, when available, an appropriate isotopic internal standard. Details of reaction conditions are provided in the *Supporting Information* and the figure legends. Quenched reaction solutions were filtered to remove protein (described in the *Supporting Information*) and analyzed on an Agilent 1260 HPLC using a SeQuant ZIC-pHILIC (5 μm, 2.1 x 100 mm) with buffer A (90% acetonitrile, 10% water, 10 mM ammonium formate, pH 4) and buffer B (10% acetonitrile, 90% water, 10 mM ammonium formate, pH 4). A linear gradient from 80% to 40% A over 16 min and held at 40% A for 1 min at a flow rate of 0.2 mL/min. Mass spectra were acquired in positive ionization mode on an Agilent 6460 QQQ. Source and acquisition parameters were: gas temperature 300 °C, drying gas 11 L/min, nebulizer 45 psi, capillary voltage 3500 V, fragmentor 60-76 V, and acquisition rate 3 spectra/s.

## RESULTS AND DISCUSSION

### Substrate-triggered formation of a 618-nm-absorbing intermediate in BesC

Whereas rapid mixing of an anoxic solution containing BesC (initially apo) and ≥ 2 molar equivalents of Fe(II) with O_2_-saturated buffer resulted only in slow development of stable ultraviolet absorption (Fig. 1A), inclusion also of L-Lys, 4-Cl-Lys (racemic at both C2 and C4), or *S*-(2-aminoethyl)-L-cysteine (4-thia-L-Lys) in the protein reactant solution led to rapid development of an intense, transient visible absorption feature centered at ∼ 618 nm (Fig. 1B, C). This feature is reminiscent of those associated with µ-peroxodiiron(III) complexes in UndA,^23^ SznF,^24^ and multiple FDOs.^17, 37–39^ Of the two previously studied HDOs, substrate triggering of intermediate formation was also seen for UndA but not for SznF. BesC lacks the extra site 1 carboxylate ligand seen in the x-ray crystal structure of Fe(II)_2_-SznF and, in accordance with the correlation noted in that study, is also triggered by substrate binding.

**Figure 1.**
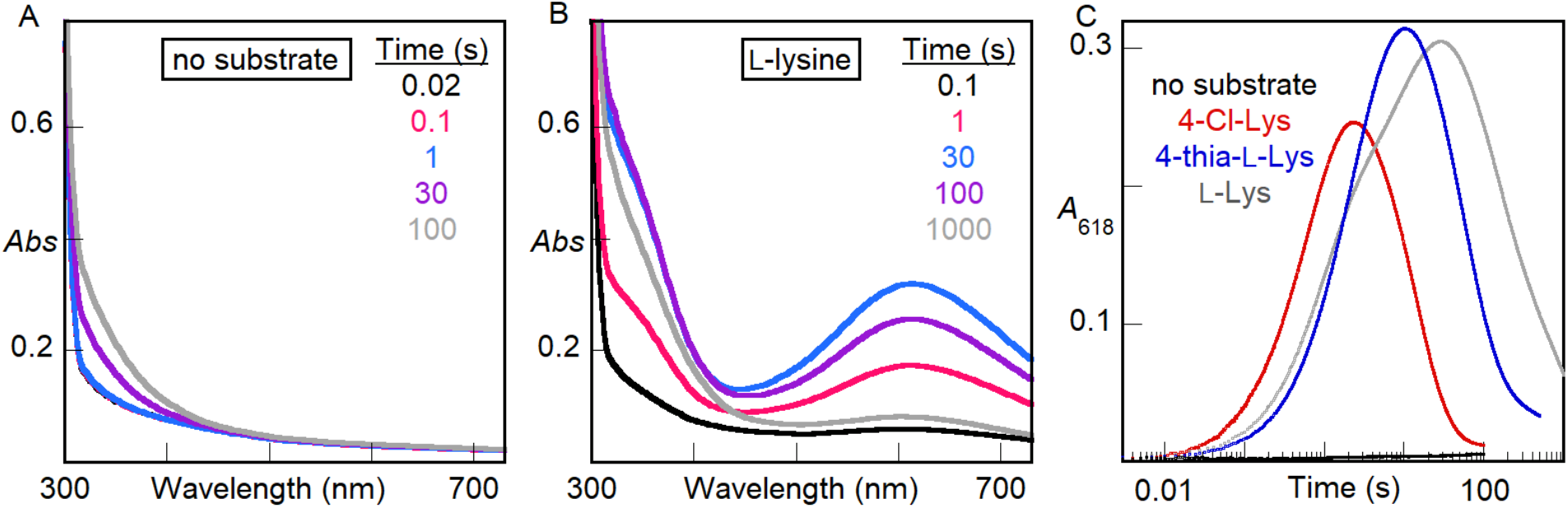
Absorption spectra acquired after rapid mixing at 5 °C of an anoxic solution of BesC (0.30 mM) and Fe(II) (0.60 mM, 2 molar equiv) in the (***A***) absence or (***B***) presence of 1 mM L-Lys with an equal volume of O_2_-saturated buffer. (***C***) Kinetic traces showing the accumulation and decay of the absorbing intermediate as a function of time in the presence of 1 mM (initial concentration) of the substrate indicated by the color-coded legend. A control lacking substrate is shown in *black*.

In the reaction with L-Lys at 5 °C, complete decay of the transient absorption feature requires tens of minutes, but in the reactions with 4-thia-L-Lys and 4-Cl-L-Lys, the decay is ∼ 5-fold and ∼ 30-fold faster, respectively (Fig. 1C). The sensitivity of the lifetime of the complex to substrate modifications at C4 (H *→* Cl or C *→* S), the site of the only C–H bond that is cleaved in the fragmentation reaction (Fig. S2, showing formation of primarily *d*_3_-L-allylglycine from *d*_4_-L-Lys), implicates the absorbing species as an authentic intermediate, a conclusion supported by the observations of primary deuterium kinetic isotope effects (D-KIE) presented below.

### Mössbauer-spectroscopic evidence that the 618-nm-absorbing species is a µ- peroxodiiron(III) complex

Mössbauer spectra – acquired at 4.2 K with a 53-mT field applied parallel to the *γ* beam – of samples prepared by freeze-quenching the BesC•Fe(II)•L-Lys/O_2_ reaction at times within the lifetime of the absorbing species confirm its assignment as a µ-peroxodiiron(III) complex. The spectrum of the frozen BesC•Fe(II)•L-Lys reactant solution (Fig. 2A, *spectrum a*) exhibits features that can be analyzed as two partially resolved quadrupole doublets with isomer shift (*δ*) and quadrupole splitting (ΔE_Q_) parameters characteristic of high-spin Fe(II) ions (*δ* ∼ 1.2 mm/s, ΔE_Q_ ∼ 3 mm/s; the actual parameters used to generate the green fit line are given in Table S3). The spectra of samples frozen between 1 s and 120 min after a mix (at 5 °C) of this solution with an equal volume of O_2_-saturated buffer exhibit diminished contributions from these quadrupole doublet features, reflecting oxidation of the Fe(II). At shorter reaction times (e.g., 1 s, Fig. 2A, *spectrum b*), a new quadrupole doublet develops (*blue lines*), reaching its maximum intensity in the sample frozen at 31 s (Fig. 2A, *spectrum c*), near the time of maximum *A*_618_ in the stopped-flow experiments. From the difference of the experimental spectra of the 1-s freeze-quench and reactant samples (Fig. 2C, *spectrum a*), the parameters of this new doublet were determined to be *δ* = 0.58 mm/s and ΔE_Q_ = 1.15 mm/s (Fig. S3). These parameters are similar to those reported for the ∼ 600-nm-absorbing, antiferromagnetically coupled, high-spin µ-peroxodiiron(III) enzyme intermediates in SznF, UndA, and FDOs (as well as synthetic models thereof), thus confirming the assignment of the BesC intermediate as such a complex.^1–2, 40–41^ Between reactions times of ∼ 30 s and ∼ 300 s, the spectrum of the intermediate partially decays, giving rise to a new quadrupole doublet (Fig. 2B, *spectrum a*). Subtractions of the spectra of the 31-s and 1-s samples (Fig. 2C, *spectrum b*) and of the 301-s and 31-s samples (Fig. 2C, *spectrum c*) resolve the features of the new doublet (*red lines*) and define its parameters as *δ* = 0.51 mm/s and ΔE_Q_ = 1.00 mm/s (Fig. S4). This spectrum is attributed to the diiron(III) product state generated upon decay of the µ-peroxodiiron(III) intermediate. At much longer reaction times (tens-hundreds of minutes), the diiron(III) complex also begins to decay, as paramagnetic features characteristic of uncoupled high-spin Fe(III) ions – either bound in mononuclear fashion within BesC or free in solution – develop. By a reaction time of 120 min, the paramagnetic features dominate the spectrum, contributing more than 60% of the total absorption area (Fig. S5). These observations establish that BesC behaves as UndA and SznF in undergoing spontaneous disintegration of its diiron(III) product cluster. As previously noted, this behavior contrasts with that of FDOs, which stably bind their diiron(III) cofactors, and emphasizes the dynamic nature and quasi-stability of the HDO architecture that contributes the two iron sites.

**Figure 2.**
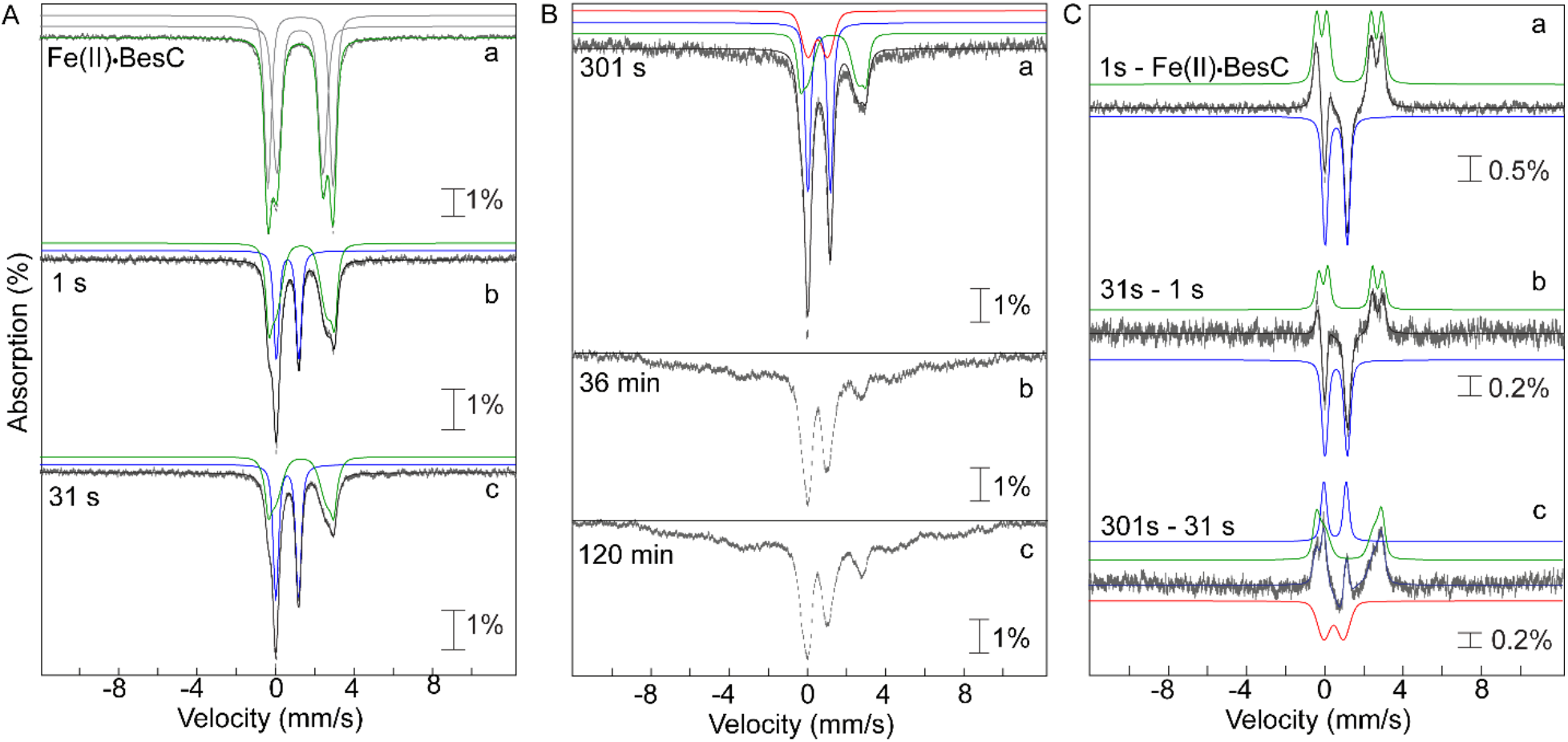
4.2 K Mössbauer spectra, acquired with a 53 mT magnetic field applied parallel to the direction of propagation of the γ-beam, of freeze-quenched samples from the reaction of Fe(II)·BesC with O_2_. The experimental spectra are depicted by grey vertical bars of heights reflection the standard deviations of the absorption values during acquisition of the spectra. The black solid lines are the overall simulated spectra, and the colored lines are theoretical spectra illustrating the fractional contributions from the Fe(II)·BesC reactant complex (green), the µ-peroxodiiron(III) intermediate (blue), and the diiron(III) product (red), to the experimental spectra. Values of the parameters and relative areas of these spectra are provided in Table S2. (***A***) Spectra of samples freeze-quenched at short reaction times, during formation and decay of the µ-peroxodiiron(III) intermediate: anoxic Fe(II)·BesC reactant complex; (*b* and *c*) spectra of samples quenched at 1 s and 31 s, respectively. (***B***) Spectra of samples frozen at longer reaction times: (*a*) 301 s, 36 min, and (*c*) 120 min. These spectra demonstrate the disintegration of the diiron(III) “product” cluster and accumulation of uncoupled high-spin Fe(III) species at longer reaction times. (***C***) Difference spectra: (*a*) 1 s − Fe(II)·BesC reactant complex; (*b*) 31 s − 1 s; and (*c*) 301 s − 31 s. These spectra illustrate the changes associated with conversion of the Fe(II)·BesC reactant to the µ-peroxodiiron(III) intermediate and its subsequent conversion to diiron(III) product.

### Kinetic evidence that the µ-peroxodiiron(III) complex or its successor abstracts H• from C4 of the substrate

Comparison of the decay kinetics from reactions with L-Lys of natural isotopic abundance and 4,4,5,5-[^2^H_4_]-L-Lys (*d*_4_-L-Lys) at two temperatures reveals a normal deuterium kinetic isotope (D-KIE) at either temperature (Fig. 3A). Regression “fits” of the averaged kinetic traces from two trials for each isotopolog at each temperature by the equation for a single exponential decay (dashed lines in Fig. 3A) are not ideal but lead to an observed D-KIE (*k*_obs_,_H_/*k*_obs,D_) of 2.1 ± 0.1 at either temperature. Fits by the equation for two parallel decay processes (solid lines in Fig. 3A) are better (especially for the data from the reactions at 22 °C) and afford similar amplitudes for the two hypothetical decay processes and D-KIEs of 3.8 (5 °C) and 2.3 (22 °C) on the more isotope-sensitive, slower phase. These values are all too large to be associated with a secondary D-KIE, implying that the intermediate is involved in HAT from C4 of the substrate, most simply as the direct H• acceptor (Scheme S1B, *black lower pathway*). Although it has been proposed in computational studies,^42^ little experimental evidence for HAT to a peroxodiiron(III) complex has been reported. The lone exception of which we are aware is the work by Lippard and co-workers showing direct reaction of the µ-peroxodiiron(III) intermediate, **H_peroxo_**, in soluble methane monooxygenase with dialkyl ether substrates,^37^ which have C–H bonds that are, by virtue of the non-bonded electrons on the central oxygen, somewhat activated relative to the C4-H bonds in L-Lys.

**Figure 3.**
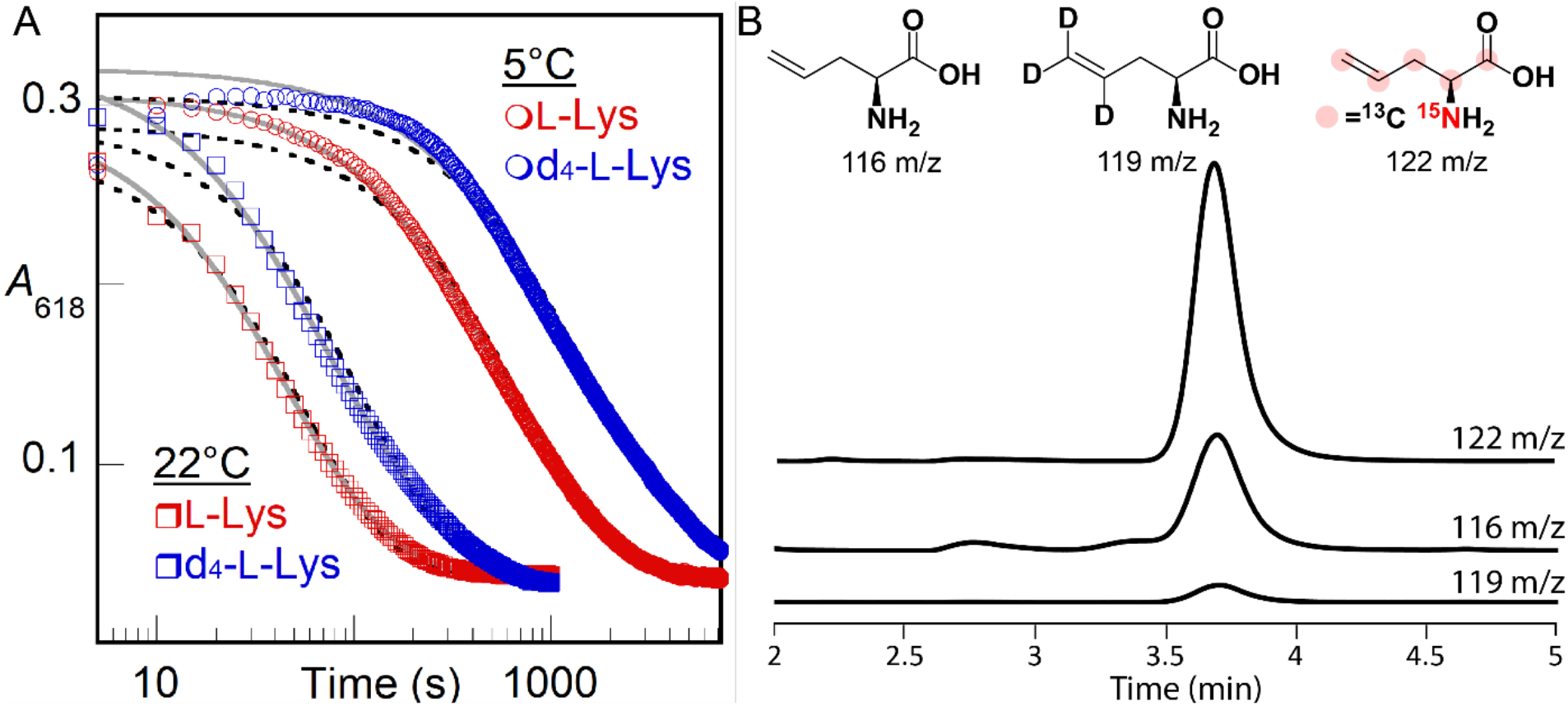
(***A***) *A*_618nm_-versus-time traces monitoring decay of the µ-peroxodiiron(III) intermediate in BesC after manual mixing of a solution containing BesC, Fe(II), and L-Lys (*red traces*) or *d*_4_-L-Lys (*blue traces*) with O_2_-saturated buffer [100 mM sodium HEPES, pH 7.5, 100mM NaCl, 5% (*v/v*) glycerol] at either 5 °C (*circles*) or 22 °C (*squares*). Final concentrations after mixing were 0.20 mM BesC, 0.40 mM Fe(II), 12 mM (*d*_4_)-L-Lys, and ∼ 0.9 mM O_2_. Fits by the equations for single exponential decay (*black dashed lines*) and two parallel decay processes *(solid gray lines*) are shown. (***B***) LC-MS detection of products in single-turnover reactions of BesC (0.90 mM) after incubation with Fe(II) (1.8 mM) and *d*_4_-L-Lys or [^13^C_6_,^15^N_2_]-L-Lys (5 mM) containing L-allylglycine standard (0.050 mM) in 100 mM sodium HEPES, pH 7.5, 100 mM NaCl, 5% (*v/v*) glycerol. Single ion monitoring (SIM) of the reaction with *d*_4_-L-Lys detected a diminished peak for *d*_3_-allylglycine at 119 *m/z* relative to the peak at 122 *m/z* for [^13^C_5_,^15^N]-L-allylglcine produced in the reaction with [^13^C_6_,^15^N_2_]-L-Lys. Peaks have been normalized to that from the L-allylglycine internal standard at 116 *m*/*z*.

The observed D-KIE, although sufficiently large to be assigned as a primary effect, is considerably less than those determined for HATs from unactivated carbon centers to oxidized iron intermediates in other enzymes.^8, 43–46^ These D-KIEs can be very large (*k*_H_/*k*_D_ > 50) as a consequence of the much greater contribution of quantum-mechanical tunneling to ^1^H transfer than to ^2^H transfer. A large intrinsic D-KIE on such a step can be kinetically masked by either (i) slow, reversible, disfavored conversion of the complex to a more reactive intermediate that is the actual H• abstractor (Scheme S2A) or (ii) decay of the absorbing complex through one or more additional, isotope-insensitive (i.e., unproductive) pathway(s) that become(s) dominant when the HAT-initiated, productive pathway is drastically slowed by a large intrinsic D-KIE (Scheme S2B).^44, 47^ To assess whether the latter mechanism of kinetic masking is operant in BesC-catalyzed L-Lys fragmentation, we compared yields of the L-allylglycine product in single-turnover reactions with protium- and deuterium-bearing L-Lys isotopologs. To enable use of commercially available compound as an internal standard for both reactions, we used a commercial heavy-atom-labeled L-Lys ([^13^C_6_,^15^N_2_]-L-Lys, with 99% isotopic enrichment in both ^13^C and ^15^N) as the protium-bearing substrate and the *d*_4_-L-Lys for comparison. Given that the magnitudes of primary ^13^C- and ^15^N-KIEs are generally less than 1.1, we anticipated that any such effects would not obscure the vastly dominant impact of deuterium substitution on the reaction efficiency, as determined by LC-MS quantification of the [^13^C_5_,^15^N]- and [4,5,5-^2^H_3_]-L-allylglycine products against the natural-isotopic-abundance standard (Fig. 3B). In six trials with each isotopolog at two different BesC concentrations [0.30 and 0.90 mM with 2 equiv Fe(II)], the [^13^C_6_,^15^N_2_]-L-Lys yielded a mean and range of 12 ± 3 times the quantity of L-allylglycine produced from 4,4,5,5-[^2^H_4_]-L-Lys, implying that the presence of deuterium severely uncouples decay of the µ-peroxodiiron(III) complex from L-allylglycine production, presumably by derailing the initiating HAT from C4 (Scheme S1B, *black lower pathway*). The observed rate constants for decay of the intermediate observed spectrophotometrically (*k*_obs,H_, *k*_obs,D_) will be the sum of the rate constants for both productive (*k*_H_ or *k*_D_) and uncoupled decay (*k*_unc_) (Eq. 1 and 2), whereas the coupling efficiencies with the two substrates (*CE*_H_ and *CE*_D_) will be

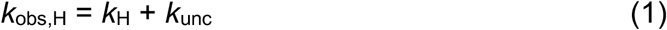

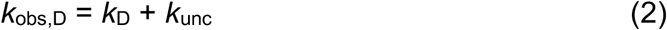

given by ratios of *k*_H_ and *k*_D_ to the sum of rate constants for coupled and uncoupled decay pathways (Eq. 3 and 4). Even without knowledge of the precise values of the coupling

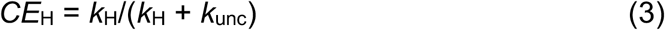

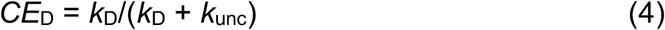

efficiencies, a system of five independent equations can be solved for all five unknown parameters (*k*_H_, *k*_D_, *k*_unc_, *CE*_H_ and *CE*_D_) using the observed rate constants for intermediate decay (*k*_obs,H_ = 0.0135 s_-1_ and *k*_obs_,_D_ = 0.0063 s^-1^ at 5 °C) and measured *ratio* of coupling efficiencies (Eq. 5) to yield *k*_H_ = 0.00075 s^-1^, *k*_D_ = 0.000031 s^-1^, *k*_unc_ = 0.0060 s_-1_, *CE*_H_ =

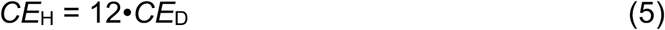

0.55 (55% coupled) and *CE*_D_ = 0.049 (4.9% coupled). The fact that decay of the intermediate in the reaction with 4-Cl-Lys is so much faster than in the reaction with L-Lys (∼ 30-fold) suggests that BesC is likely to be more faithful in coupling O_2_ activation with fragmentation of its native, chlorine-bearing substrate. Regardless, the analysis yields a resolved D-KIE (*k*_H_/*k*_D_) of 24 for production of L-allyglycine, which should be dominated by the primary effect on H/DAT from C4 to the µ-peroxodiiron(III) intermediate or the hypothetical successor with which it reversibly interconverts.

A secondary D-KIE on C5-C6 fragmentation could, theoretically, divert some of the flux in the reaction of the *d*_4_-L-Lys to a different product (e.g., 4-hydroxy-L-Lys). This normal secondary effect, which should be ≤ 1.8, would inflate the value – presumed above to originate from a primary D-KIE on C4 HAT – estimated on the basis of L-allylglycine yields. The conservative assumptions of (1) a normal secondary D-KIE as large as 2, causing 2-fold less L-allyglycine to be produced in favor of an alternative product, and (2) a negligible heavy-atom (^13^C, ^15^N) primary KIE on L-allylglycine production from the [^13^C_6_,^15^N_2_]-L-Lys substrate (which, if not negligible, would oppose the effect of the secondary D-KIE) would lead to an estimate of ∼ 6 for the impact of (specifically) the primary D-KIE on the ratio of coupling yields, *CE*_H_/*CE*_D_. Using this estimate in place of Eq 5 above would result in a calculated *k*_H_/*k*_D_ of ∼ 13. The conclusion that the true primary D-KIE is actually greater than the semi-classical limit of 7-8 makes it more plausible that the µ-peroxodiiron(III) complex could indeed be the H•-abstracting complex in the fragmentation of L-Lys.

### Evidence that 4-thia-L-Lys can undergo OAT to S4

The conclusion that the µ-peroxodiiron(III) complex or its successor cleaves the C4-H bond of the substrate implies that 4-thia-L-Lys should not be fragmented but should rather undergo a different reaction, with the most likely possibility being OAT to the sulfur, as has been shown to occur in other iron enzymes that naturally oxidize sulfur-containing substrates^48–50^ or are challenged with sulfur-containing substrate analogs.^51^ We tested by LC-MS for possible fragmentation products from 4-thia-L-Lys (*S*-methyl-L-cysteine, L-cysteine, L-serine, L-alanine, glycine) but did not detect such products. We readily detected a species with *m*/*z* of 181, +16 relative to the M-H^+^ ion of 4-thia-L-Lys, only in the complete reaction; reactions from which BesC, Fe(II) or O_2_ was omitted did not produce the associated species (Fig 4). Use of ^18^O_2_ caused the new LCMS peak to shift to *m*/*z* = 183, +18 relative to the M-H^+^ ion of 4-thia-L-Lys, showing that the substrate undergoes oxygenation. Because all feasible carbon hydroxylations in this compound would produce unstable hemiaminal (C2 or C6) or thiohemiacetal (C3 or C5) species that would readily fragment in water, the stable incorporation of an oxygen atom into the detected product necessarily implies that an OAT to a heteroatom has occurred. Although OAT to the *ε*-amine could, in principle, occur upon derailment of an L-Lys fragmentation pathway initiated by amine radical formation, the presence of sulfur in place of C4 would be expected to *promote* rather than to impede C5-C6 cleavage by this mechanism, owing to its well-known ability to stabilize an *α*-carbon-centered radical, in this case on C5. OAT to the sulfur is by far the more likely outcome and is consistent with the observed primary D-KIE on decay of the intermediate.

**Figure 4.**
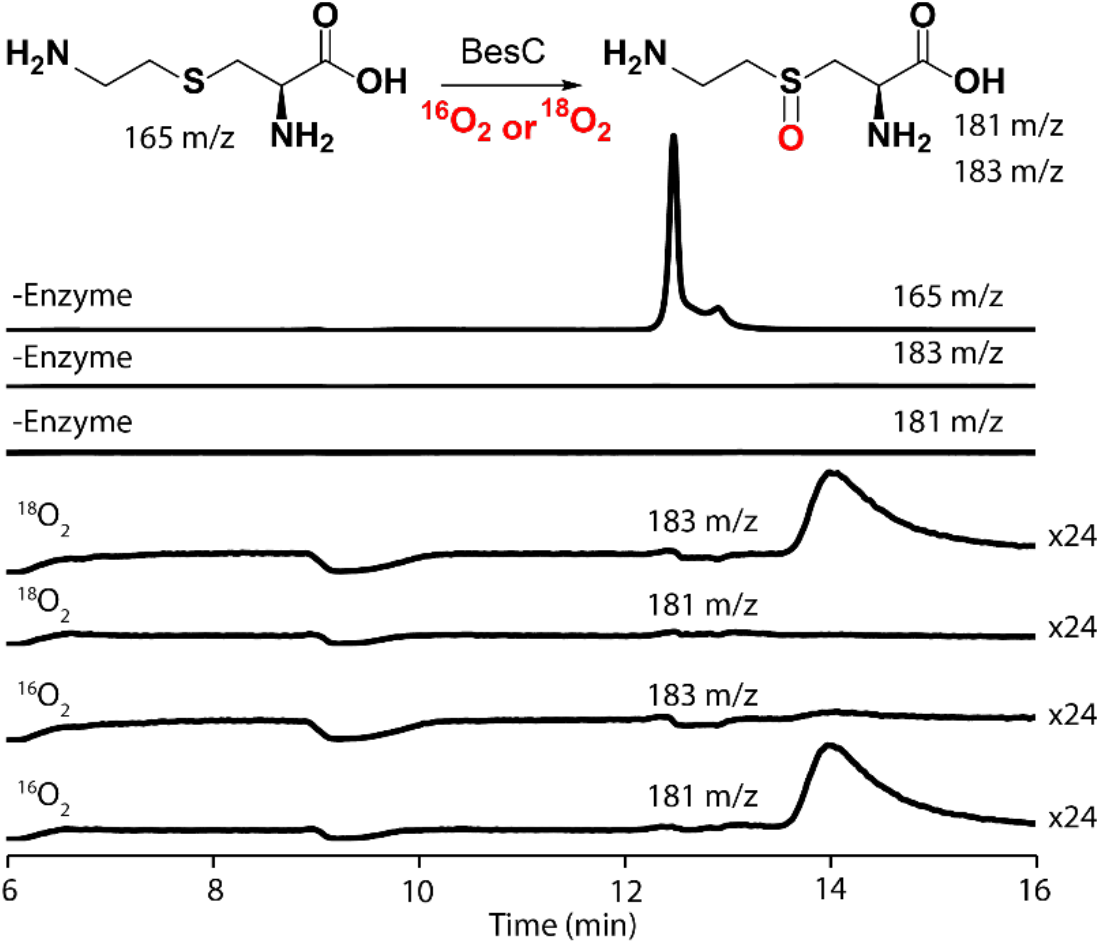
LC-MS detection of products in single-turnover reactions of BesC (0.25 mM) after incubation at 22 °C with Fe(II) (0.50 mM) and 4-thia-Lys (0.25 mM) in 100 mM sodium HEPES, pH 7.5, 100 mM NaCl, 5% *(v/v)* glycerol. Single-ion monitoring (SIM) of a control reaction without enzyme shows a large peak for 4-thia-L-Lys at 165 *m/z* (*rear trace*) but no peaks at either 183 or 181 *m*/*z*. In the presence of natural abundance O_2_ (^16^O_2_), the substrate is mostly consumed, and SIM traces at 181 *m/z* reveal a new peak with a shift of +16 *m/z* compared to 4-thia-Lys. Reactions with ^18^O_2_ show a mass shift of +18 *m/z* (at 183 *m/z*), consistent with the incorporation of an oxygen atom into the product.

### Structural determinants of substrate triggering

We qualitatively explored the determinants of substrate triggering by comparing the capacities of a series of analogs to promote intermediate accumulation at a single fixed concentration (1 mM before mixing) (Fig. S6). BesC is surprisingly promiscuous in this trait: L-lysine analogs modified by stereoinversion of the *α*-carbon (D-Lys), addition or deletion of a side chain methylene (L-homolysine or L-ornithine), desaturation between existing methylenes (*trans*-4,5-dehydro-DL-lysine) replacement of the *ε*-amine by hydroxyl or hydrogen [(6-hydroxy)-L-norleucine (6-OH-L-Nle) or L-norleucine], or mono- or di-methylation of N7 all support accumulation of the peroxide complex to varying extents under these conditions (Fig. S6). By contrast, trimethylation of N7 or replacement of the *α*-carboxylate or *α*-amine by hydrogen (1,6-diaminohexane, 6-aminocaproic acid) abolishes intermediate accumulation. The data suggest that, despite cleaving the side chain of (4-chloro)-L-Lys, BesC relies more on the *α*-carbon substituents as binding determinants, tolerating even a likely cation-to-polar substitution of N7 by O. The latter observation hints at the possibility that the active site could promote N7 deprotonation, which is expected to be required for either radical or polar fragmentation following HAT from C4 (Scheme. S1B).

As groundwork for establishing the physical basis of substrate triggering in BesC, we defined the concentration dependencies (Fig. 5) for L-Lys (***A***), 6-OH-Nle (***B***), and 4-thia-L-Lys (***C***) with Fe(II) fixed at 2 molar equivalents relative to BesC. With sub-saturating substrate, the *A*_618_ kinetic traces, reflecting formation and decay of the µ-peroxodiiron(III) complex, exhibit two distinct formation phases. We attribute the faster phase to the capture of O_2_ by the properly pre-assembled BesC•Fe(II)_2_•substrate complex and the slower phase to association of a missing component [Fe(II) or substrate] followed by reaction with O_2_. Fitting the traces to the analytical solution for absorbance in a system of parallel processes with two different, transparent reactants, A and A’, but a common absorbing intermediate, B, and transparent product, C (Scheme S3), which approximates the two-step reaction affording the slower formation phase as a single step to simplify analysis), allows the amplitudes of the fast phase and thus the relative concentrations of BesC•Fe(II)_2_•substrate complex in the reactant solutions to be extracted. Regression fits of plots of this quantity versus the concentration of the substrate yield approximate dissociation constants (*K*_D_) of 0.3 ± 0.1 mM for L-Lys, 10 ± 1 mM for 6-OH-L-Nle, and < 30 μM for 4-thia-L-Lys (Fig. 5 insets). The traces from the 4-thia-L-Lys reaction are not as clearly biphasic, and the effect of the analog saturates at just slightly greater than one molar equivalent, implying that the association is tight and this analog is particularly effective at driving assembly of the O_2_-reactive complex (Fig. 5C). These results are consistent with a binding selectivity for a Lys analog with a larger, more polarizable substituent at the 4 position, as in 4-Cl-L-Lys.

**Figure 5.**
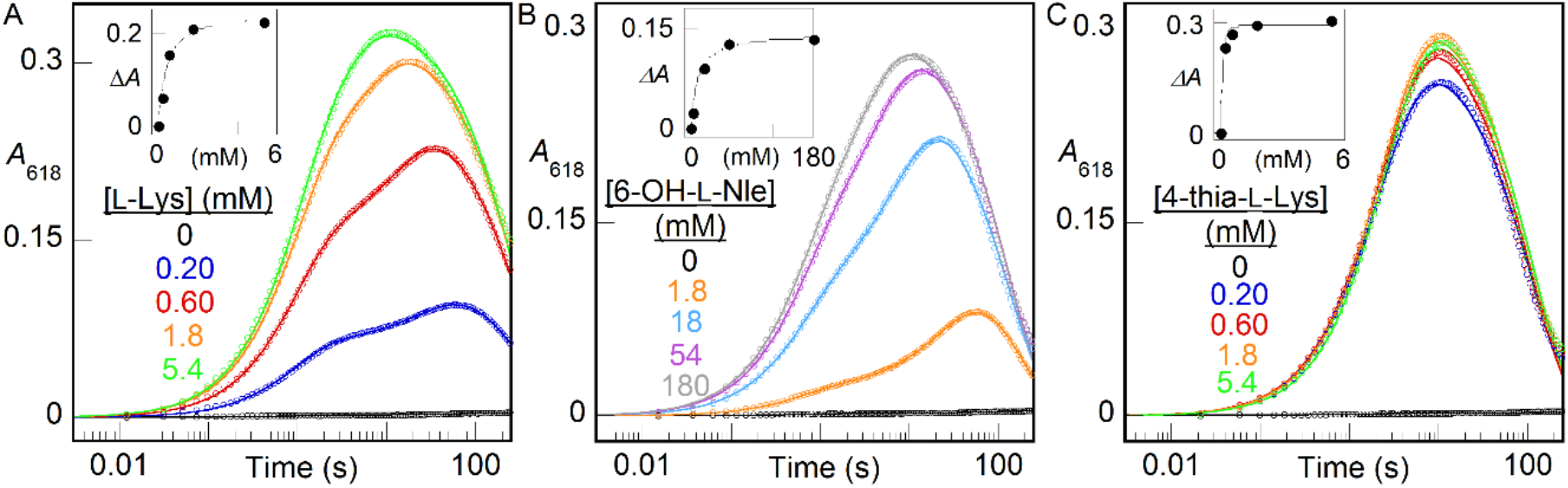
*A*_618_-versus-time traces demonstrating the effect of substrates on triggering the reaction of Fe(II)•BesC complex with O_2_ at 5°C. A reactant solution of BesC (0.20 mM), Fe(II) (0.40 mM, 2 molar equiv), and (***A***) L-Lys, (***B***) 6-OH-L-Nle, or (***C***) 4-thia-L-Lys at the concentrations indicated by the color-coded legends was mixed with an equal volume of O_2_-saturated buffer. The insets plot the amplitude of the fast phase (Δ*A*) as a function of the substrate concentration. The solid lines are fits of the quadratic (*A* and *C*) or hyperbolic (*C*) equation for binding to these data.

### Physical basis of substrate triggering: synergy of substrate and Fe(II) binding

Given published evidence that the HDO scaffold generally affords only one stable Fe(II) site (at most) and at least one site (site 2) that is improperly configured (i.e., intrinsically disordered) prior to a conformational change driven by metal binding,^19,23,25–26,52^ we considered that substrate triggering in BesC could reflect synergy in substrate and Fe(II) binding, perhaps involving substrate-driven configuration of the dynamic site 2. As a first assessment, we titrated the triggering effect of L-Lys with 1.0 (Fig. 6A), 2.0 (Fig. 5A) or 6.0 (Fig. 6B) molar equivalents of Fe(II) in the BesC solution. All three plots show the same qualitative behavior, including a hyperbolically increasing fast formation phase with increasing [L-Lys]. With 1.0 equiv Fe(II), the transient feature saturates at less than 50% of its maximum possible amplitude with excess Fe(II) (Fig 6B), consistent with the fact that this quantity of Fe(II) is only half that needed to fill both sites of the cofactor. Linkage between Fe(II) and L-Lys binding is evident by comparing traces from reactions with 2 and 6 equiv of Fe(II) at sub-saturating concentrations of L-Lys (e.g., 0.20 mM and 0.60 mM): the greater Fe(II) concentration yields considerably greater amplitude in the fast phase (Fig. 6B, *solid traces*) than the lesser [Fe(II)] (*dotted traces*), a difference that is largely overcome at the highest [L-Lys]. Conversely, the effect of increasing [Fe(II)] on the fast-phase amplitude is much more pronounced with 0.20 mM L-Lys (Fig. 6C) than with 5.4 mM L-Lys (Fig. 6D). These effects of varied [L-Lys] at fixed [Fe(II)] and varied [Fe(II)] at fixed L-Lys reflect, generally, synergistic metal and substrate binding.

**Figure 6.**
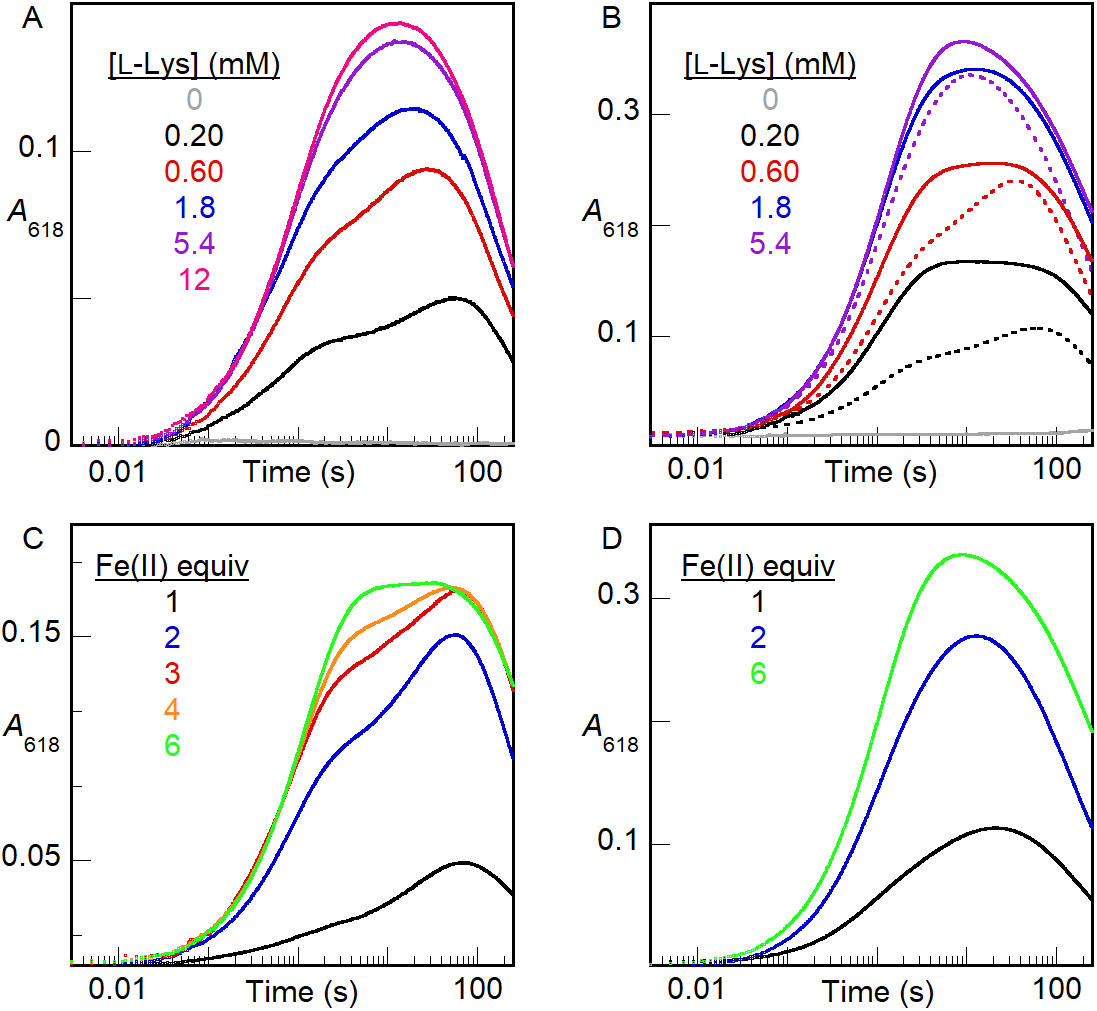
*A*_618_-versus-time traces from stopped-flow absorption experiments in which an anoxic solution of BesC (0.20 mM), Fe(II) and L-Lys was mixed at 5 °C with an equal volume of O_2_-saturated buffer. In titrations of L-Lys, BesC was incubated with (***A***) 0.20 mM (1 molar equiv) or (***B***) 1.2 mM (6 molar equiv) of Fe(II). The dotted traces in panel B are from matching experiments (from Figure 5A) with 0.40 mM (2 molar equiv) Fe(II) and are overlaid to illustrate the impact of varying [Fe(II)].

We sought more direct evidence of this linkage by using a kinetic assay based on the Fe(II) indicator, ferrozine, which forms the purple Fe(II)•ferrozine3 complex with *λ*_max_ = 562 nm, and *ε*_562_ = 27,900 M^-1^cm^-1^. Fe(II)_aq_ is complexed rapidly (∼ 50 s^-1^ under the conditions of the assay), whereas Fe(II) bound by BesC must dissociate before it can be detected._53-54_ Mixing an anoxic solution containing BesC and 0.8-1.0 equiv Fe(II) with an anoxic solution containing excess ferrozine leads to development of absorbance at 562 nm (*A*_562_) with a half-life (*t*1_/2_) of 0.5-1 s (longer at higher [BesC], owing to rebinding by the protein in competition with ferrozine capture^54^). No discernible fast phase of development is seen, implying that BesC can bind one Fe(II) ion tightly in the absence of a substrate (Fig. 7A, *black trace*, and Fig. S7). With 2.0 equiv Fe(II), phases with approximate *t*_1/2_ values of ∼ 0.02 s, arising from Fe(II)_aq_, and ∼ 1 s, arising from Fe(II) bound by BesC, have approximately equal amplitudes, suggesting that BesC binds *only one equiv Fe(II)* under these conditions (Fig. 7B, *black trace*). Indeed, inclusion of three times as much Fe(II) (6 equiv) increases the amplitude only of the fast phase arising from chelation of Fe(II)_aq_ (Fig. S7); the amplitude of the slow phase from release/chelation of Fe(II) bound in BesC remains essentially constant, establishing that binding of a second Fe(II) ion is, at best, much weaker than binding of the first Fe(II) ion. Inclusion of 12 mM L-Lys (Fig. 7A, *blue*) or 4-thia-L-Lys (*red*) in the BesC solution with 0.8 equiv Fe(II) slows dissociation of the complex, extending the half-life of *A*_562_ development to ∼ 6 s or ∼ 40 s, respectively. This slowing of ferrozine chelation reflects some combination of a diminished rate of dissociation from, and increased rate of binding by, BesC in the presence of bound substrate. Slower dissociation would, obviously, slow colorimetric detection, and faster association could cause released Fe(II) to partition more frequently toward rebinding by BesC in competition with irreversible capture in the low-spin Fe(II)•ferrozine_3_ complex. Again, the greater affinity of the C4 *→* S analog is evident, in this case by its greater delay of Fe(II) release and detection. More strikingly, the presence of this high concentration of either substrate in the BesC reactant solution containing 2.0 equiv Fe(II) drastically suppresses the fast phase of complexation (Fig. 7B, *blue and red*), almost completely abolishing it for the case of the tighter-binding 4-thia-L-Lys (*red*). The amplitude of the slow phase associated with bound Fe(II) increases to account for the loss from the fast phase. These results establish that substrate binding allows BesC to complete assembly of its functional cofactor by binding the second Fe(II) ion. Given the aforementioned observations in prior structural studies on SznF, the most likely explanation for this phenomenon is that the substrate induces a conformational change that properly configures an otherwise intrinsically disordered site 2.

**Figure 7.**
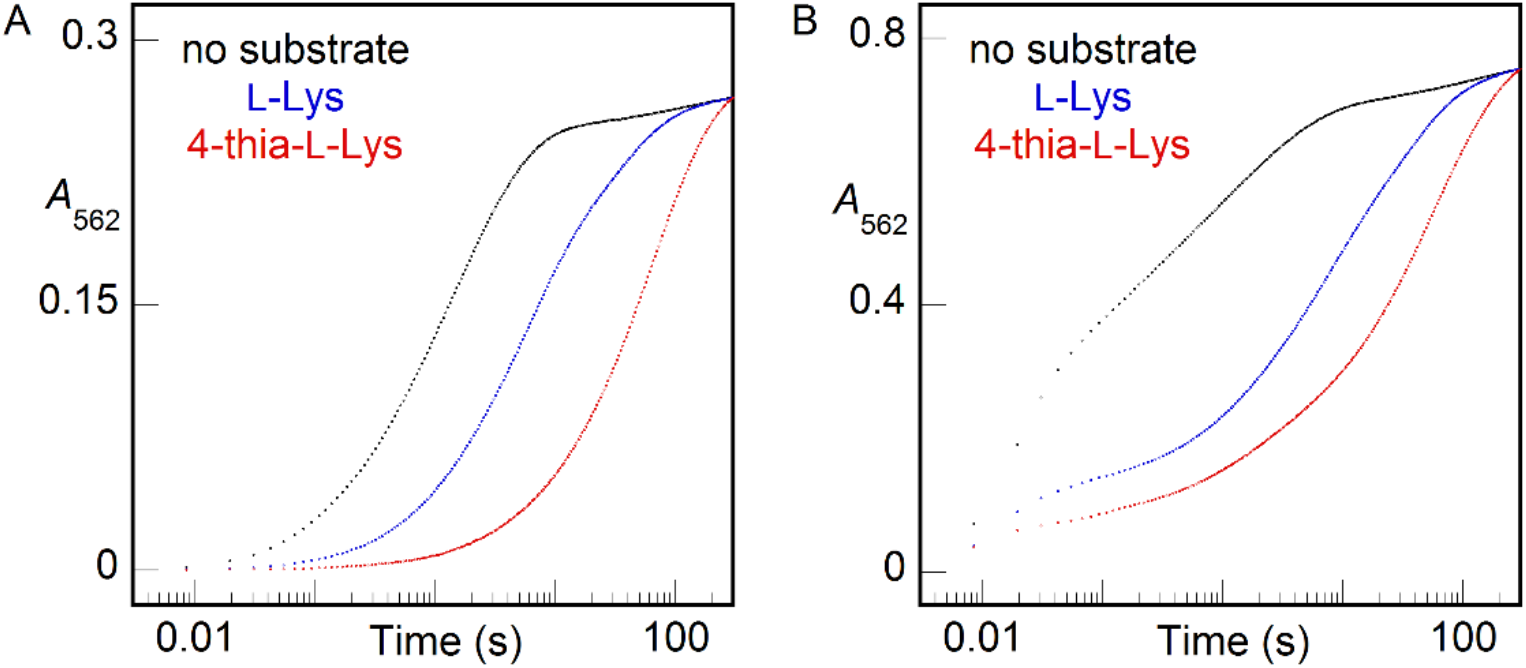
*A*_562_-versus-time traces monitoring chelation of Fe(II) by ferrozine, rapidly when free in solution or more slowly after its rate-limiting dissociation from BesC. An anoxic solution of 0.16 mM apo BesC was loaded with either (***A***) 0.13 mM (0.8 molar equiv) or (***B***) 0.32 mM (2 molar equiv) Fe(II) in the absence substrate (*black*) or in the presence of 12 mM L-Lys (*blue*) or 4-thia-L-Lys (*red*). This solution was subsequently mixed with an equal volume of an anoxic solution of 4 mM ferrozine. The absorbance of the purple Fe(II)•ferrozine_3_ complex at 562 nm (*A*_562_) was monitored as a function of reaction time. The traces shown here are representative of at least 2 trials for each condition. Results of experiments with greater Fe(II):BesC ratios in the absence of substrate are provided in Fig. S7.

### Substrate dissociation from, and binding to, the µ-peroxodiiron(III) intermediate

Given the proposed mechanism by which L-Lys triggers intermediate formation (properly configuring an otherwise disordered Fe site 2), its relatively modest affinity, and the sluggishness of its reaction with the intermediate (or its successor), we wondered if the addition of O_2_ to the cofactor effectively traps the substrate on the enzyme or, alternatively, if it can dissociate after driving configuration of the second iron site and formation of the intermediate. Protection of reactive intermediates is a well-known phenomenon in enzymology, and conformational changes that effectively seal the active site are often seen or invoked. For the case of iron-oxygen adducts, there is ample evidence that protonation occurs in forward conversion to more reactive states or as part of proton-coupled reduction steps (including HAT), and so sequestration from solvent and sources of protons would seem necessary to contain a reactive complex for the tens of minutes seen in the case of the µ-peroxodiiron(III) complex formed in BesC with L-Lys. Conversely, dissociation/binding of substrate from/to such a state would seem incompatible with prolonged stability. Surprisingly, L-Lys does indeed dissociate from the long-lived intermediate to allow for binding and reaction of another substrate. We demonstrated this behavior by sequential-mix stopped-flow experiments (Fig. 8), in which the intermediate was allowed to accumulate maximally following an initial mix of BesC•Fe(II)_2_•L-Lys complex with O_2_-containing buffer and this solution was then mixed with an excess of one of the more reactive substrates, 4-thia-L-Lys (*blue*) or 4-Cl-Lys (*red*). Both substrates caused concentration-dependent (Fig. S8) acceleration of decay of the intermediate relative to controls in which the pre-formed intermediate was mixed with buffer or additional L-Lys. The fact that neither substrate reacted with the intermediate as rapidly as it does in a single-mix experiment, in which it triggers intermediate formation itself, suggests that L-Lys dissociation is slower than the chemical reactions of the two more reactive substrates with the intermediate. Nevertheless, the fact that substrate can dissociate and re-bind at all in the intermediate state again underscores the dynamic character of the HDO scaffold and its potential to be leveraged in biocatalysis applications.

**Figure 8.**
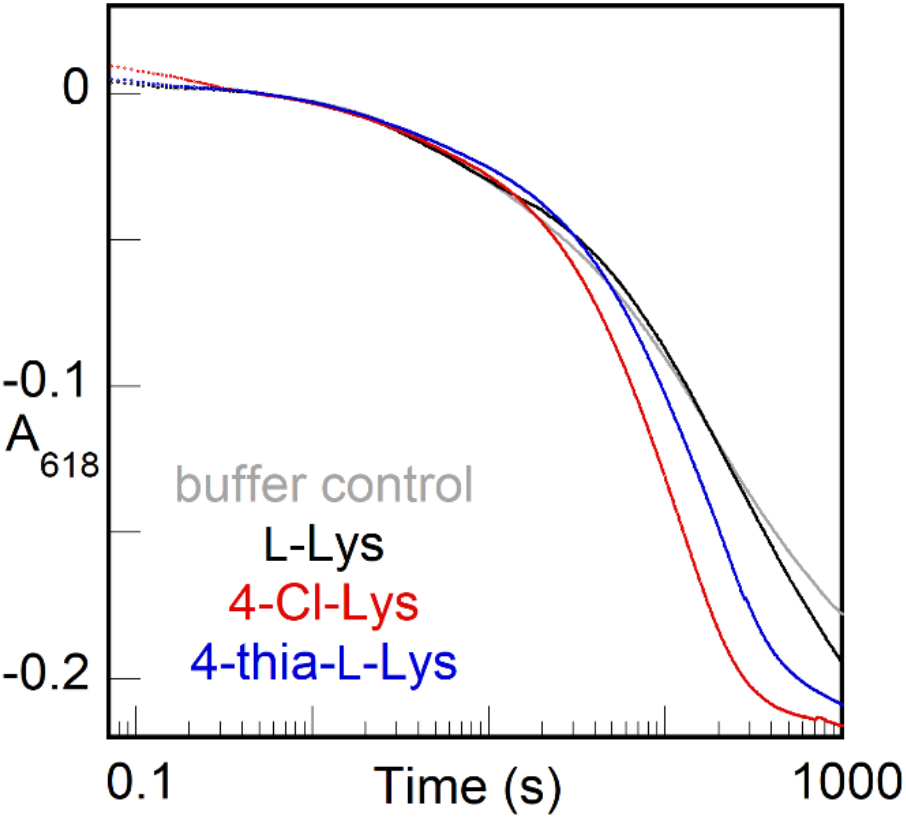
*A*618-versus-time traces from a sequential-mix stopped-flow experiment showing accelerated of decay of the BesC µ-peroxodiiron(III) intermediate when mixed with a more reactive substrate after triggering of formation by L-Lys. An anoxic solution containing BesC (0.40 mM), Fe(II) (0.80 mM), and L-Lys (2.7 mM) was mixed with an equal volume of O2 saturated buffer at 5 °C. After aging for 30 s, this solution was mixed with an equal volume of an anoxic solution of buffer (*gray*), excess L-Lys (10 mM, *black*), 4-Cl-Lys (6.8 mM, *red*), or 4-thia-L-Lys (6.8 mM, *blue*).

## CONCLUSIONS

The members of the emerging HDO family studied to date (i) all bind iron only weakly in (at least) one of two sites, (ii) form accumulating µ-peroxodiiron(III) intermediates, and (iii) allow or promote disintegration of the diiron(III) cofactor product following intermediate decay. Structural studies have suggested that core helix *α*3, which provides three cofactor ligands to iron 2, is generally dynamic and responsible for the weak binding of Fe(II) at this site, the need for a conformational change prior to Fe(II) binding, and the instability of the oxidized cofactor. BesC shares these emerging, unifying traits. Its promiscuous triggering by a soluble substrate and its variously modified derivatives enabled demonstration that the triggering event involves substrate-promoted configuration of an initially disordered second iron site, most likely site 2, to allow assembly of the active BesC•Fe(II)_2_•substrate and capture of O_2_. The resultant µ-peroxodiiron(III) complex is relatively long-lived despite the ability of the triggering substrate to dissociate and reassociate in the intermediate state, and reactivity trends as well as a primary D-KIE on decay of the intermediate of ∼ 13-24 in the reaction with L-Lys show that the intermediate or a successor with which it interconverts initiates the complex fragmentation by HAT from C4. As the most likely successor to the observed intermediate would be a diiron(IV) complex with cleaved O-O bond, the results presented herein would provide either (1) just the second example (of which we are aware) of experimental evidence for HAT directly to a peroxodiiron(III) intermediate or (2) the first example (of which we are aware) of reversible O-O-bond cleavage in such a complex.

## ACCESSION NUMBERS

*S. cattleya* BesC F8JJ25 (Uniprot ID)

## ACKNOWLEDGMENT

This work was supported by the National Science Foundation (CHE-1610676 to C.K., J.M.B., and A.K.B. and CHE-1710588 to M.C.Y.C) and the National Institutes of Health (GM138580 to J.M.B., GM119707 to A.K.B., GM127079 to C.K.). J.W.S acknowledges support of the National Institute of General Medical Sciences of the National Institute of Health (F32GM136156). M.E.N. acknowledges the support of a National Science Foundation Graduate Research Fellowship. The content is solely the responsibility of the authors and does not necessarily represent the official views of the National Institute of Health.

## AUTHOR INFORMATION

Corresponding Authors *

## Notes

The authors declare no competing financial interest.

