## Supplementary information for "A Substrate-triggered µ-Peroxodiiron(III) Intermediate in the 4-Chloro-L-Lysine-Fragmenting Heme-Oxygenase-like Diiron Oxidase (HDO) BesC: Substrate Dissociation from, and C4 Targeting by, the Intermediate"

### SUPPLEMENTARY PROCEDURES

*Reactions of the Fe(II)•BesC substrate complex with $O_2$ (Figure 1).* Reactant solutions contained 0.3 mM BesC, 0.6 mM (2 molar equiv) Fe(II), and 1 mM of substrate/analog in 100 mM sodium HEPES (pH 7.5), 100 mM NaCl, and 5% glycerol (*v/v*). In control samples that lacked substrate, the compound was omitted from the initial reaction mixture. The anoxic protein solution was mixed at 5 °C with an equal volume of the same buffer at equilibrium (at 5 °C) with ~ 1.05 atm $O_2$ (~1.8 mM). Absorbance was monitored with the photodiode array detector.

*Freeze-Quench Mössbauer (FQ-Möss) Spectroscopy (Figure 2).* Freeze-quench Mössbauer samples were prepared according to previously published procedures.[1] The BesC•Fe(II)$_2$•L-Lys reactant complex was assembled from anoxic solutions of 1.7 mM BesC, 3.4 mM $^{57}$Fe(II), and 5.1 mM L-Lys. An $O_2$-stable, acidic stock solution of $^{57}$Fe(II) was prepared from commercial $^{57}Fe^0$, as previously described.[2] In the anoxic chamber, this solution was diluted to 50 mM Fe(II) by mixing with 1 M sodium HEPES, pH 7.5. An appropriate volume of this dilute stock solution was added to an anoxic solution of apo BesC in 100 mM sodium HEPES buffer (pH 7.5), 100 mM NaCl, 5% (*v/v*) glycerol to yield the reactant solution described above. This sample was mixed at 5 °C with an equal volume of the same buffer that had been saturated with $O_2$ (~ 1.8 mM). This solution was allowed to incubate for the varying reaction times indicated in Fig. 2 and subsequently frozen by injection into cold (-150 °C) 2-methylbutane (for reaction times from milliseconds to tens of seconds) or by pipetting into a Mössbauer cell cooled on a metal block that was in contact with liquid $N_2$ (for reaction times of minutes to hours).

*BesC Single-Turnover Assays with 4,4,5,5-[$^2H_4$]-L-lysine or [$^{13}C_6$,$^{15}N_2$]-L-lysine (Figure 3B)*. Reactions were carried out in the MBraun anoxic chamber in sealed LC-MS vials with pierceable septa. A 25 μL aliquot of a solution containing BesC (0.60 or 1.8 mM) in 100 mM sodium HEPES, pH 7.5, 100 mM NaCl, 5% glycerol, $(NH_4)_2FeSO_4$ (1.2 or 3.6 mM, 2 molar equiv), and substrate (10 mM) was mixed with an equal volume $O_2$-saturated buffer. After 2 h at 5 °C, reactions were quenched with 1.5 volumes (75 μL) of methanol and 1% formic acid containing 83 μM L-allylglycine. Control reactions omitted one of the reaction components. Samples were centrifuged at 10,000*g* for 10 min to remove precipitated protein. The supernatant was filtered by centrifugation in a 10 kDa Nanosep filter (PALL Corporation) at 10,000*g* for 10 min. Samples were analyzed by LC-MS as described in the main text.

*BesC Single-Turnover Assays with 4-thia-L-Lys (Figure 4)*. Reactions were carried out in a MBraun anoxic chamber in sealed LC-MS vials with pierceable septa. A 25 μL aliquot of a solution containing BesC (0.50 mM) in 100 mM sodium HEPES, pH 7.5, 100 mM NaCl, 5% glycerol, $(NH_4)_2FeSO_4$ (1 mM or 2 molar equiv), and 4-thia-L-Lys (0.50 mM, 1 molar equiv) was mixed with equal volume of $O_2$-saturated buffer containing either natural abundance $O_2$ or $^{18}O_2$. After 30 min, reactions were quenched with 1.5 volumes (75 μL) of methanol and 1% formic acid containing 50 μM L-norleucine. Control reactions omitted one of the reaction components. Samples were centrifuged at 10,000*g* for 10 min to remove precipitated protein. The supernatant was filtered by centrifugation in a 10 kDa Nanosep filter (PALL Corporation) at 10,000*g* for 10 min. Samples were analyzed by LC-MS, as described in the main text.

*Titrations of the Fe(II)•BesC substrate complex.* Reactant solutions contained 0.2 mM BesC; 0.2, 0.4 or 1.2 mM (1, 2 or 6 molar equiv) Fe(II); and 0, 0.1, 0.2, 0.6, 1.8, 5.8 12, 18, 54, or 180 mM substrate/analog. These samples were prepared in 100 mM sodium HEPES (pH 7.5), 100 mM NaCl, and 5% glycerol (*v/v*). The anoxic protein solution was mixed at 5 °C with an equal volume of the same buffer at equilibrium (at 5 °C) with ~ 1.05 atm $O_2$ (~1.8 mM). Absorbance was monitored with the photodiode array detector.

*Titrations of the BesC•L-lysine complex with Fe(II).* Reactant solutions contained 0.2 mM BesC; 0.2 mM, 5.4 mM, or 12 mM (1, 27, or 60 molar equiv) L-Lys or 4-thia-L-Lys; and 0.2, 0.4, 0.6, 0.8, or 1.2 mM Fe(II). These samples were prepared in 100 mM sodium HEPES (pH 7.5), 100 mM NaCl, and 5% glycerol (*v/v*). The anoxic protein solution was mixed at 5 °C with an equal volume of the same buffer at equilibrium (at 5 °C) with ~ 1.05 atm $O_2$ (~1.8 mM). Absorbance was monitored with the photodiode array detector

*Reaction of the Fe(II)•BesC complex with ferrozine.* The SF lines were soaked overnight in a solution of 5 mM sodium dithionite prepared anoxically in 100 mM sodium HEPES (pH 7.5), 100 mM NaCl, 5% glycerol (*v/v*). For the experiment, a solution containing 160 μM BesC was incubated with 50 μM sodium dithionite in 100 mM sodium HEPES (pH 7.5), 100 mM NaCl, 5% glycerol (*v/v*) to scrub out residual $O_2$ from the samples. To this solution, Fe(II) was added to 0.13 or 0.32 mM (0.8 or 2 molar equiv). In trials containing excess substrate/analog, 12 mM of either L-Lys or 4-thia-L-Lys was included in the protein solution. This solution was then mixed with an equal volume of an anoxic solution containing 4 mM ferrozine and 50 μM sodium dithionite in the same buffer. In samples containing half the amount of protein used, 80 uM BesC was incubated with 1 mM sodium dithionite in 100 mM sodium HEPES (pH 7.5), 100 mM NaCl, 5% glycerol (*v/v*). To this solution Fe(II) was added to a concentration of 40, 80, 120, 160, or 480 uM (0.50, 1.0, 1.5, 2.0 or 6.0 molar equiv). In trials containing excess substrate/analog, 12 mM of either L-Lys or 4-thia-L-Lys was added to the protein mixture. This solution was mixed with an equal volume of an anoxic solution containing 4 mM ferrozine and 1 mM sodium dithionite in the same buffer. In both cases, absorbance at 562 nm ($A_{562}$) was monitored at 5 °C with the photomultiplier tube using a 0.2 cm pathlength. When appropriate (in experiments with equal Fe(II) concentrations), a very slow loss of ~ 5-10% of the absorption amplitude over the course of all-day experiments was corrected for by normalization of traces to the same final amplitude.

*Reaction of the peroxodiiron(III) complex triggered by L-lysine with more reactive substrate analogs.* In a sequential mixing configuration, a reactant solution containing 0.4 mM BesC, 0.8 mM (2 molar equiv) Fe(II), and 2.7 mM L-Lys in 100 mM sodium HEPES (pH 7.5), 100 mM NaCl, and 5% glycerol (*v/v*). This anoxic protein solution was mixed at 5 °C with an equal volume of the same buffer at equilibrium (at 5 °C) with ~ 1.05 atm $O_2$ (~1.8 mM). After being allowed to react to accumulate the peroxodiiron(III) intermediate, this reaction solution was then mixed with an equal volume of either 10 mM L-Lys (control) or 0.1, 1.35, of 6.75 mM of either 4-thia-L-Lys or 4-Cl -Lys. Absorbance was monitored with the photodiode array detector.

**Scheme S1.** Mechanistic possibilities for BesC-mediated $O_2$-dependent fragmentation of (4-Cl)-L-Lys. (***A***) Pathway initiated by ε-amine oxidation. (***B***) Pathways mediated by C4 HAT.

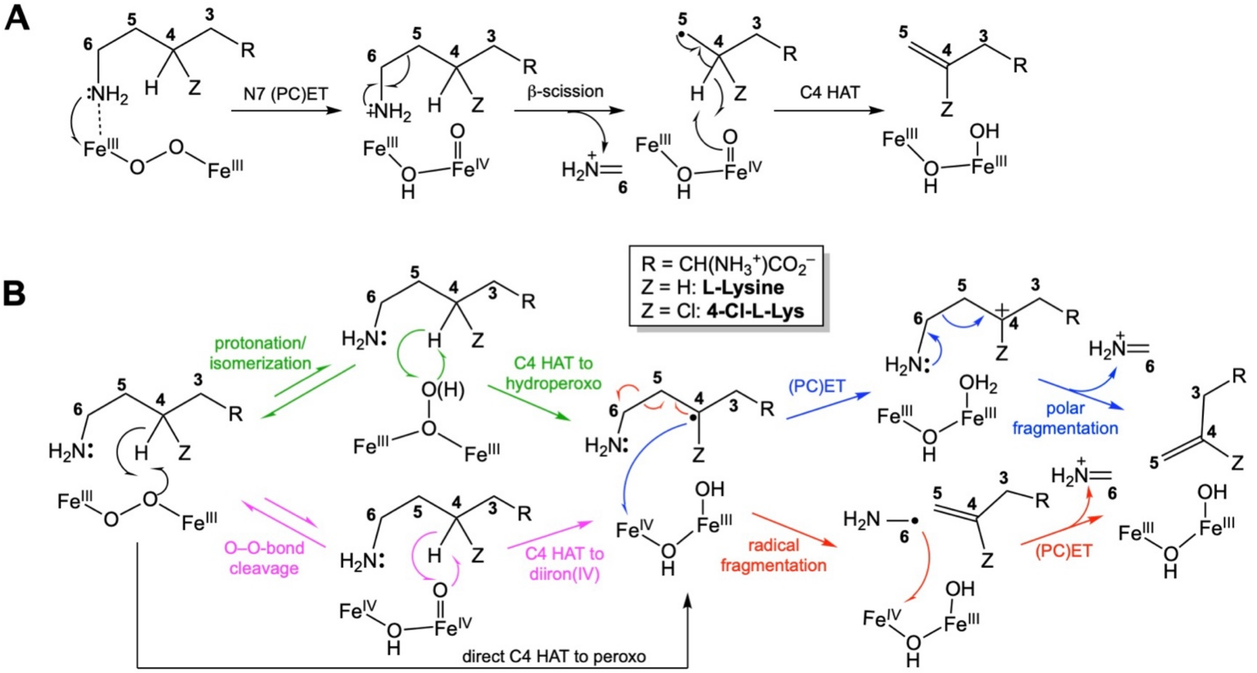

**Scheme S2**. Potential kinetic masking [in direct monitoring of μ-peroxodiiron(III) intermediate decay] of a large intrinsic D-KIE on C4 HAT by reversible conversion of the μ-peroxodiiron(III) complex to a more reactive intermediate or decay of the absorbing intermediate complex through isotope-insensitive pathway(s).

A

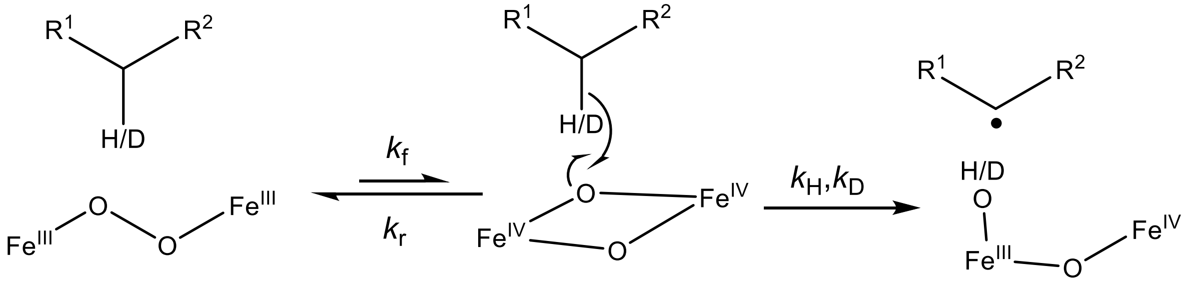


$$k_{H,obs}/k_{D,obs} = \frac{k_f \bullet k_H/(k_H + k_r)}{k_f \bullet k_D/(k_D + k_r)} = \frac{k_H/(k_H + k_r)}{k_D/(k_D + k_r)}$$

B

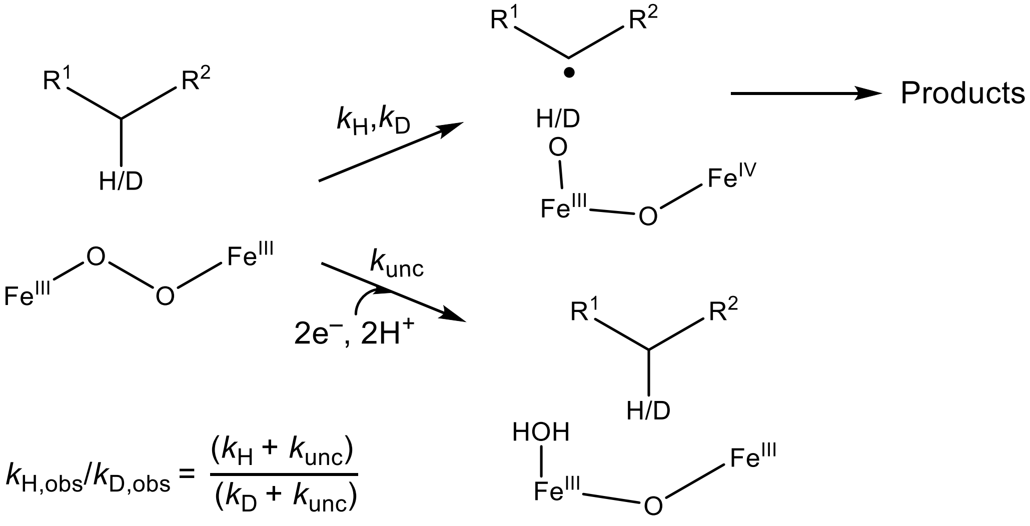


$$k_{H,obs}/k_{D,obs} = \frac{(k_H + k_{unc})}{(k_D + k_{unc})}$$

**Scheme S3**. Kinetic model used in regression analysis of the SF-Abs traces.

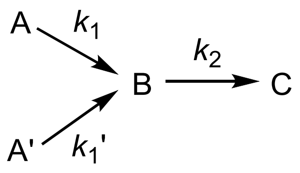

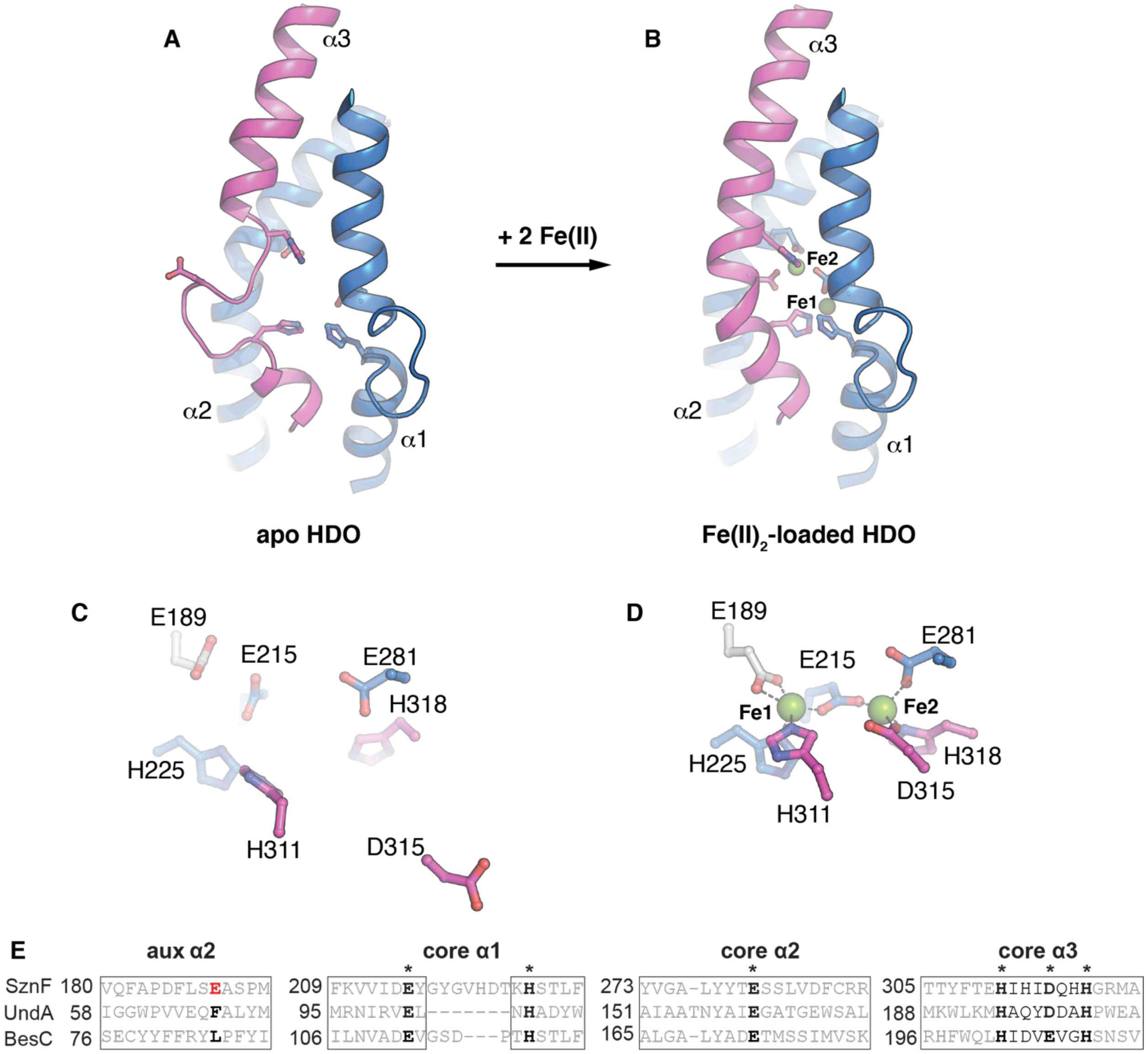


**Figure S1.** Metal loading in HDOs involves a conformational change of a core metal-binding helix. Comparison of (***A***) apo and (***B***) Fe(II)-loaded SznF reveals that core metal binding α-helix, α3 (pink), containing three protein-derived iron ligands, undergoes a significant secondary structure transition upon cofactor assembly. This change causes site 2 metal ligand, D315, to shift >10 Å from its posiiton in the apo state (***C***) to coordinate Fe2 in the fully assembled structure (***D***). A sequence alignment (***E***) shows that BesC conserves key metal binding residues in core HDO secondary structures α1-α3. BesC does not conserve the additional carboxylate ligand (*white sticks*) in an auxiiliary motif, consistent with functional assignment as a desaturase-lyase.

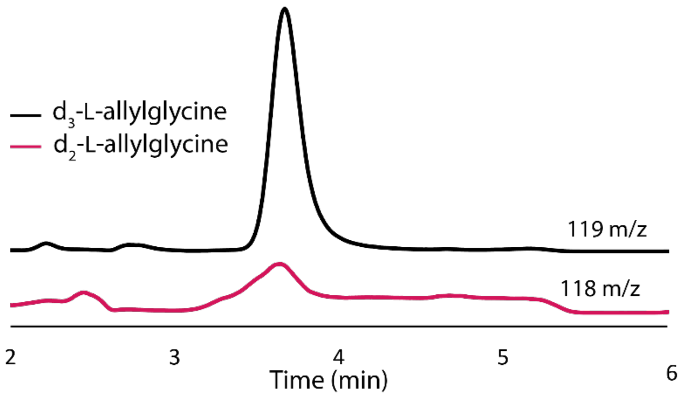


**Figure S2.** LC-MS detection of products in multiple-turnover assays of BesC (0.15 mM) after incubation with Fe(II) (0.30 mM) and $d_4$-4,4,5,5-L-Lys (1 mM) in 100 mM sodium HEPES pH 7.5, 100 mM NaCl, 5% glycerol (v/v) at 22 °C. Single ion monitoring (SIM) of the reaction with $d_4$-L-Lys shows a substantially larger peak for $d_3$-L-allylglycine at 119 *m/z* relative to the peak at 118 *m/z* that corresponds to $d_2$-L-allylglycine. The latter peak arises from a small fraction of incompletely deuterium-labeled substrate in the commercial product.

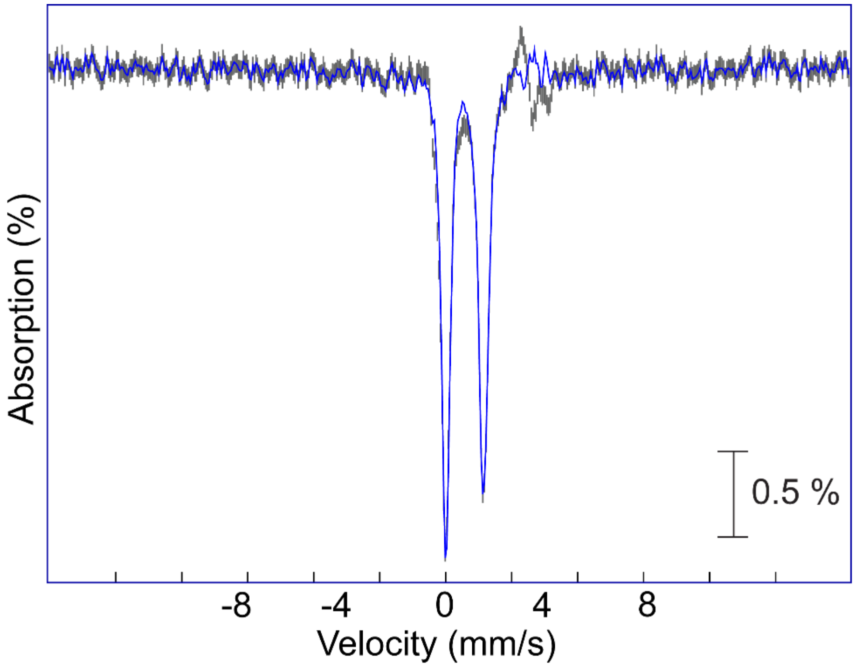


**Figure S3**. Reference spectrum of the μ-peroxodiiron(III) intermediate generated by subtraction of 57% of the experimental spectrum of Fe(II)•BesC (*grey vertical bars*) or 59% of the theoretical spectrum (*blue line*) from the spectrum of the 1-s freeze-quenched sample shown in Figure 2 of the main text.

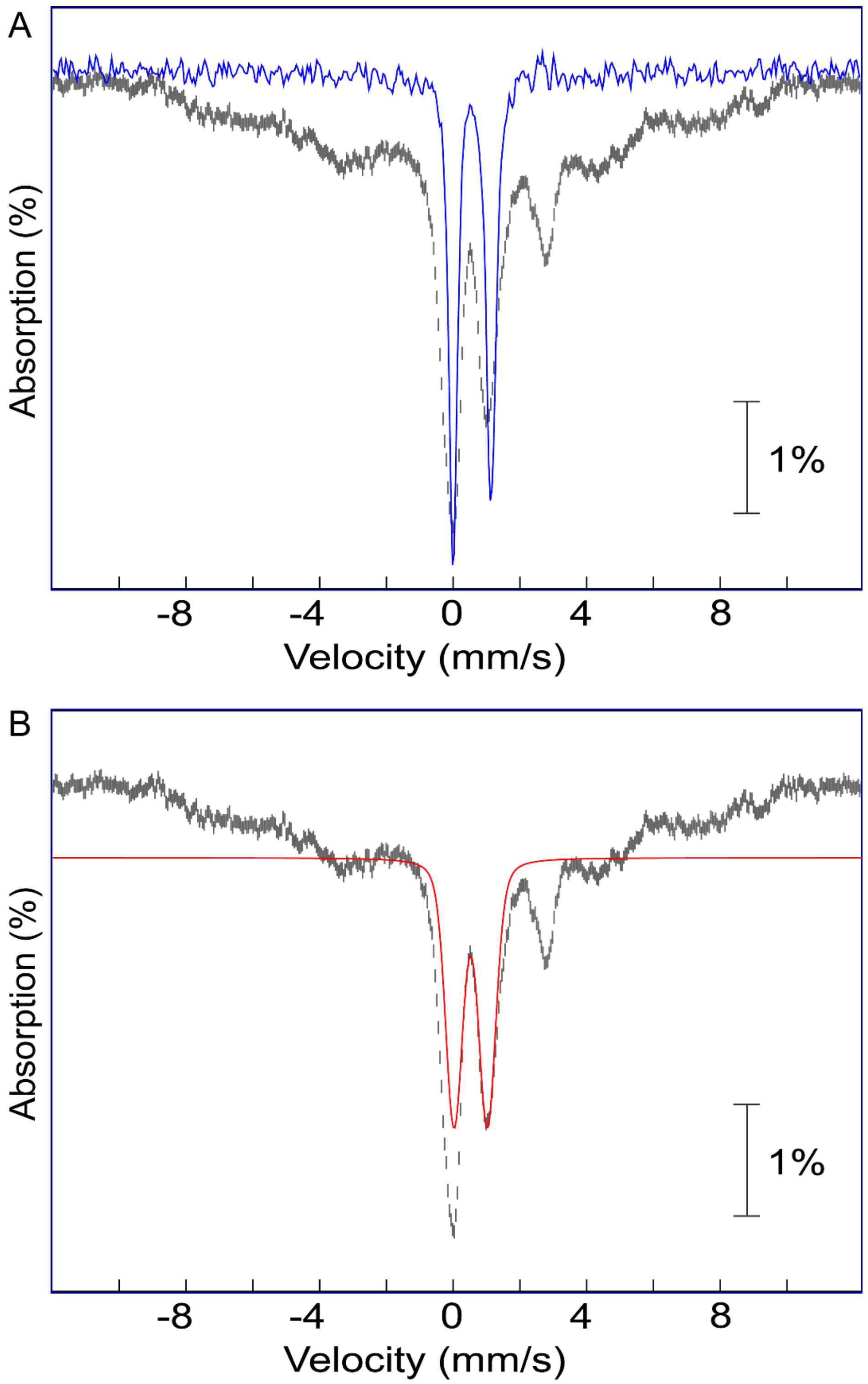


**Figure S4**. Overlay of the 4.2-K/53-mT Mössbauer spectrum with (***A***) the derived reference spectrum of the μ-peroxodiiron(III) intermediate or (***B***) the theoretical reference spectrum of the diiron(III) product.

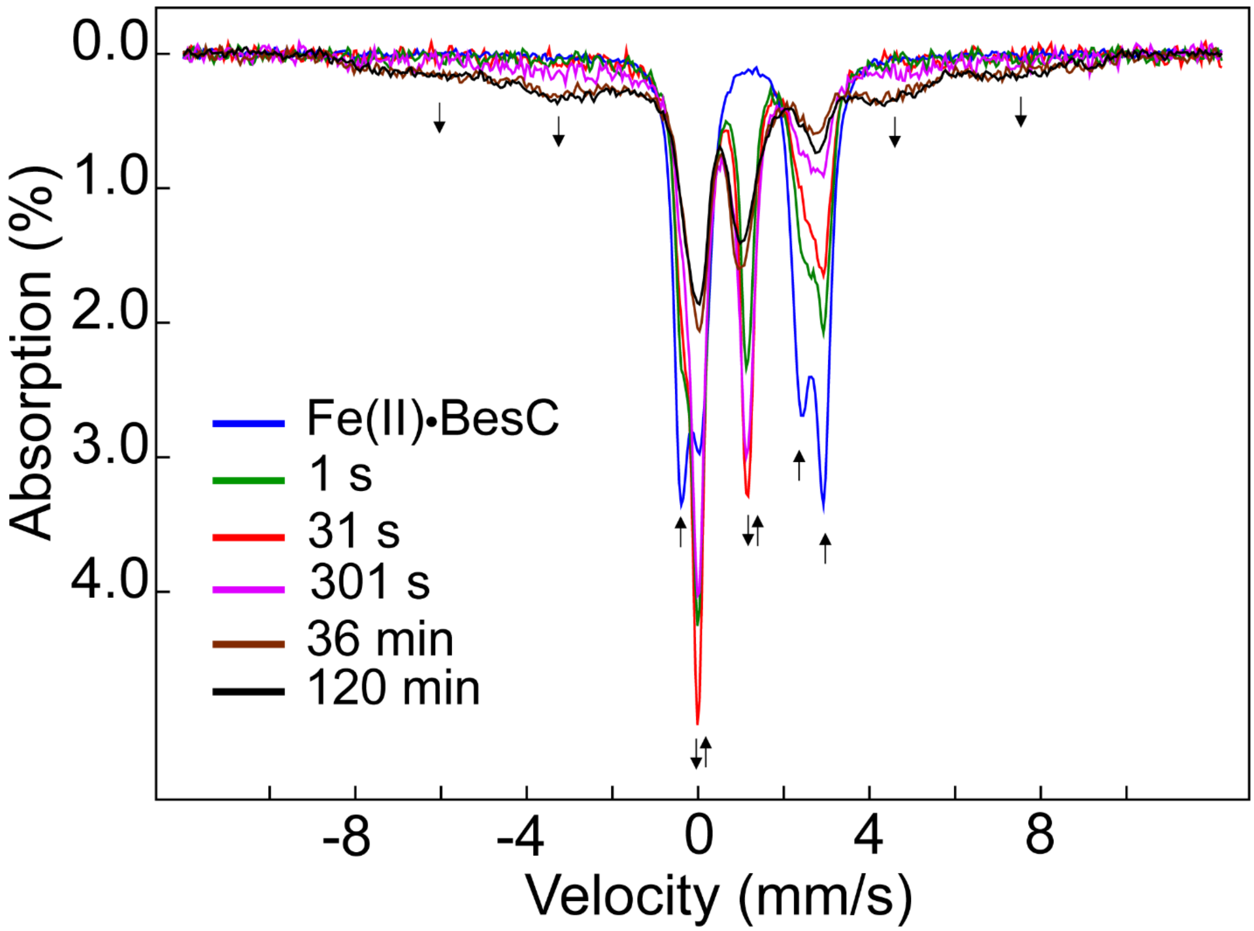


**Figure S5**. Overlay of all 4.2-K/53-mT Mössbauer spectra from the experiment depicted in Figure 2 of the main text to illustrate the development of the broad features characteristic of uncoupled high-spin Fe(III) species at long reaction times.

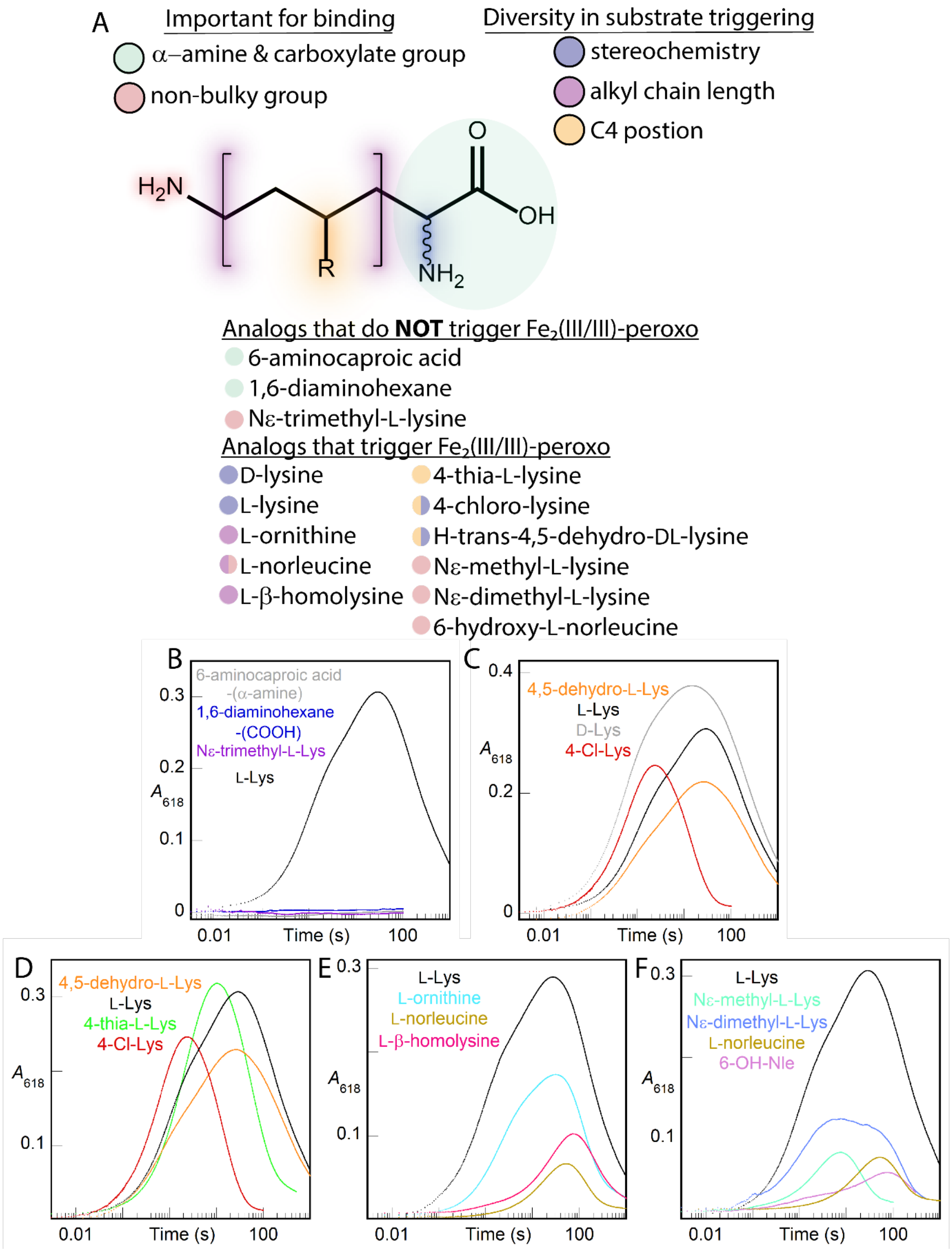
A
Important for binding
α–amine & carboxylate group
non-bulky group
Diversity in substrate triggering
stereochemistry
alkyl chain length
C4 postion
H2N
R
NH2
O
OH
Analogs that do NOT trigger Fe2(III/III)-peroxo
6-aminocaproic acid
1,6-diaminohexane
Nε-trimethyl-L-lysine
Analogs that trigger Fe2(III/III)-peroxo
D-lysine
L-lysine
L-ornithine
L-norleucine
L-β-homolysine
4-thia-L-lysine
4-chloro-lysine
H-trans-4,5-dehydro-DL-lysine
Nε-methyl-L-lysine
Nε-dimethyl-L-lysine
6-hydroxy-L-norleucine
B
6-aminocaproic acid
-(α-amine)
1,6-diaminohexane
-(COOH)
Nε-trimethyl-L-Lys
L-Lys
A618
0.3
0.2
0.1
0
0.01
Time (s)
100
C
4,5-dehydro-L-Lys
L-Lys
D-Lys
4-Cl-Lys
A618
0.4
0.2
0
0.01
Time (s)
100
D
4,5-dehydro-L-Lys
L-Lys
4-thia-L-Lys
4-Cl-Lys
A618
0.3
0.1
0.01
Time (s)
100
E
L-Lys
L-ornithine
L-norleucine
L-β-homolysine
A618
0.3
0.1
0.01
Time (s)
100
F
L-Lys
Nε-methyl-L-Lys
Nε-dimethyl-L-Lys
L-norleucine
6-OH-Nle
A618
0.3
0.1
0.01
Time (s)
100

**Figure S6**. Analysis of L-Lys analogs reveals the determinants for substrate triggering in formation of the μ-peroxodiiron(III) intermediate. (***A***) Schematic depiction of components of lysine modified in these experiments. These include (*i*) loss of the α-amine or carboxylate functional groups, (*ii*) mono- or di-methylation of the ε-amine, (*iii*) stereoinversion of the C2 position, (*iv*) change in length of the alkyl chain, and (v) subsitution at the C4 position. $A_{618}$-versus-time traces are shown for each of these five groups. All reactions were carried out by mixing an anoxic solution of BesC (0.30 mM), Fe(II) (0.60 mM) and the substrate analog (1 mM) with $O_2$-saturated buffer [100 mM sodium HEPES pH 7.5, 100 mM NaCl, 5% (v/v) glycerol] at 5°C. All plots show the kinetic trace for L-Lys for comparison. (***B***) Substrate analogs lacking the α-amine/carboxylate groups or trimethylated at the ε-amine do not trigger intermediate formation in BesC. We posit that these analogs either fail to bind in the active site (6-aminocaproic acid, 1,6-diaminohexane) or contain altered charge that prevents reactivity with $O_2$ ($N^{\varepsilon}$-trimethyl-L-Lys). (***C***) Substrate analogs with altered stereochemistry at C2 are tolerated by BesC. In the case of D-Lys, the BesC intermediate forms faster and accumulates to a greater extent. (***D***) BesC also tolerates substitutions at the C4 position. In particular, with 4-thia-L-Lys, the μ-peroxodiiron(III) intermediate decays more quickly (by a factor of 5) than with L-Lys. (***E***) The BesC intermediate is triggered by amino acids with shorter alkyl chains; however, the kinetics are markedly perturbed. (***F***) Lastly, substitutions at the ε-amine are permitted by BesC, with the exception of the trimethylated side chain. Although these substitutions are tolerated, the kinetics are similar to substrates that vary in alkyl chain length.

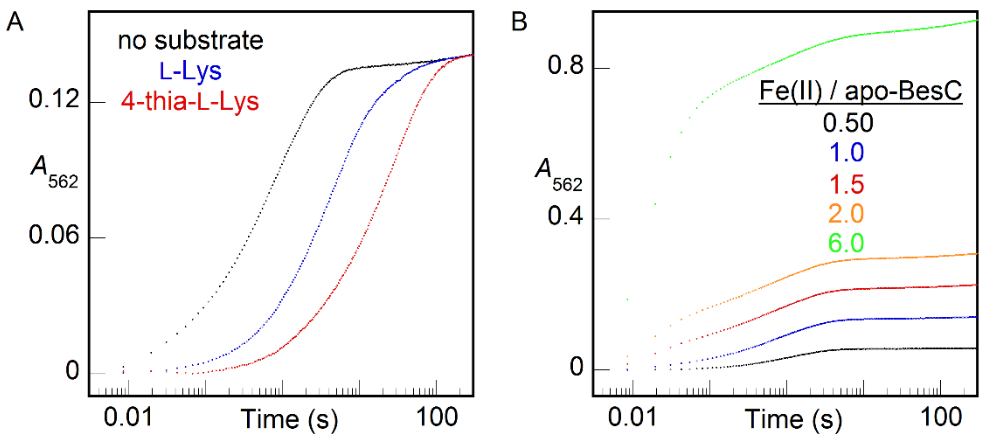


**Figure S7.** $A_{562}$-versus-time traces for the detection of Fe(II) by ferrozine following its dissociation from BesC. An anoxic solution of 80 μM apo-BesC was loaded with (***A***) 80 μM (1 molar equiv) Fe(II) in the absence of any substrate (*black*) or in the presence of 12 mM L-Lys (*blue*) or 4-thia-L-Lys (*red*); or (**B**) 40-480 μM (0.5-6 molar equiv) Fe(II) in the absence of any substrate. This solution was then mixed with an equal volume of an anoxic solution of 4 mM ferrozine, and the absorbance of the purple Fe(II)•ferrozine$_3$ complex at 562 nm ($A_{562}$) was monitored as a function of reaction time. The traces shown here are representative of at least 2 trials for each condition.

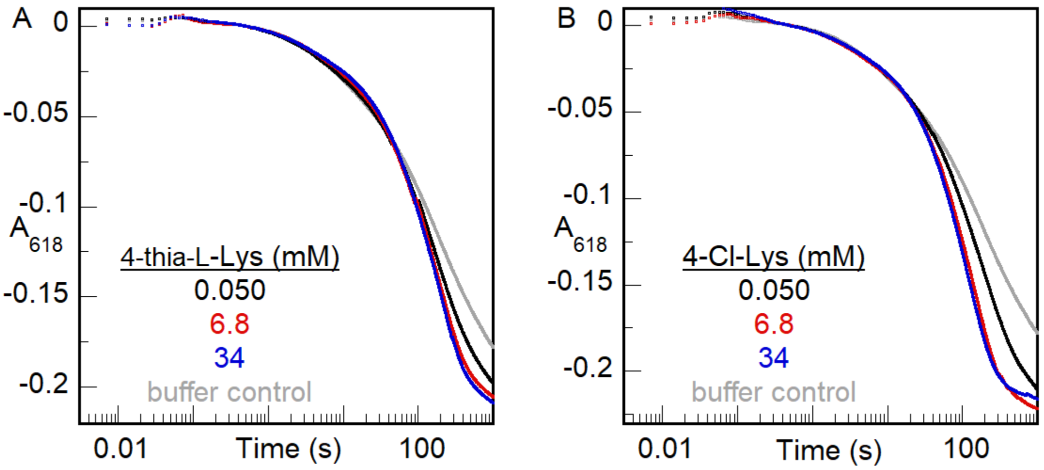


**Figure S8**. $A_{618nm}$-versus-time traces from a sequential-mix stopped-flow experiment showing decay of the μ-peroxodiiron(III) intermediate. A reaction solution containing BesC (0.40 mM), Fe(II) (0.8 mM), and L-Lys (2.7 mM) was mixed with $O_2$-saturated buffer at 5 °C for 30 s. This solution was subsequently mixed with (***A***) 4-thia-L-Lys (0, 0.050, 6.8, 34 mM), or (***B***) 4-Cl-Lys (0, 0.050, 6.8, 34 mM).

**Table S1**. Cloning primers for the preparation of BesC.

| Primer | Sequence | Restriction Enzyme Site |
|---|---|---|
| Forward | 5'- catatgaccgacctgaacaccc-3' | *NdeI* |
| Reverse | 5'- ctcgagtcacttgccgatgc-3' | *XhoI* |

**Table S2**. Recipe for 10,000X trace metals cocktail for minimal (M9) media supplementation.

| Reagent | mg/L | Final concentration in medium (nM) |
|---|---|---|
| $(NH_4)Mo_7O_{24}$ | 37.1 | 3 |
| $CoCl_2 \bullet 6H_2O$ | 71.4 | 300 |
| $H_3BO_3$ | 247 | 400 |
| $CuSO_4 \bullet 5H_2O$ | 25 | 10 |
| $MnCl_2 \bullet 4H_2O$ | 198 | 80 |
| $ZnSO_4 \bullet 7H_2O$ | 28.8 | 10 |

**Table S3.** Parameters used to simulate Mössbauer spectra.

| Fe species | δ (mm/s) | $\Delta E_Q$ (mm/s) | Linewidth (mm/s) | Relative area (%) |
|---|---|---|---|---|
| Peroxo-$Fe_2$(III/III) intermediate | 0.58 | 1.15 | -0.25 | Variable |
| $Fe_2$(III/III) successor complex | 0.51 | 1.00 | -0.80 | Variable |
| $Fe_2$(II/II) in Figure 3Aa * | 1.24 | 2.35 | -0.49 | 50 |
| | 1.27 | 3.32 | -0.373 | 50 |
| $Fe_2$(II/II) in Figure 3A, spectra b and c * | 1.27 | 2.67 | -0.84 | Variable |
| | 1.30 | 3.38 | -0.22 | variable |

*The contribution of the reactant $Fe_2$(II/II) complex(es) varies as a function of the reaction time, as is evident by the different shape of the upward-pointing peaks associated with the reactant $Fe_2$(II/II) at ~+2.7 mm/s (e.g., compare spectra a, b, and c in Figures 3 A & C in main manuscript). This observation suggests heterogeneity of the reactant. We simulated the spectrum of the $Fe_2$(II/II) complex(es) by assuming two symmetrical quadrupole doublets.
